# Benchmarking foundation models for resting-state functional magnetic resonance imaging time series in cognitive aging research

**DOI:** 10.64898/2026.11.05.764644

**Authors:** Yang Merik Liu, Meishan Ai, Kanhao Zhao, Yu Zhang, Feng Vankee-Lin

## Abstract

Foundation models trained on large neuroimaging datasets could help smaller studies investigate cognition from brain recordings. However, whether the foundation model-derived representations improve assessment of current cognition, prediction of follow-up cognition, or characterization of cognitive change remains unclear. We evaluated nine configurations from five model families that process regional resting-state functional magnetic resonance imaging time series, using 3,103 scans from 1,653 participants across two observational and five intervention aging cohorts. Primary frozen representations showed no consistent superiority over conventional functional connectivity for concurrent cognitive associations or diagnostic classification. Changing the readout improved selected concurrent cognitive associations without updating model weights, indicating that performance depended partly on how representations were extracted. Additional post-training and embedding alignment yielded gains in selected settings but did not consistently improve performance across outcomes. In the primary longitudinal analyses, imaging representations did not establish added value beyond baseline clinical measures. These findings show that readout choice influences concurrent cognitive associations, while the added value of these representations for longitudinal outcomes remains unestablished.

## Introduction

Understanding individual differences in brain function and changes within individuals can help explain cognitive aging and inform efforts to preserve cognition. Resting-state functional MRI (rs-fMRI) measures spontaneous fluctuations in the blood oxygen level-dependent (BOLD) signal, allowing brain function to be characterized without requiring participants to perform a cognitive task. Observational studies use rs-fMRI to examine differences across age and cognitive or clinical states, while intervention studies examine brain changes accompanying cognitive training, exercise, and other interventions.^1–3^ A common representation used in these studies is whole-scan functional connectivity (FC), typically estimated using correlations between regional BOLD signals, which summarizes their average statistical relationships but does not preserve temporal ordering.^4,5^ Features derived directly from BOLD time series may additionally capture temporal dependencies within and between regions and the occurrence and transitions of recurring network states.^5,6^ Previous studies have linked measures of brain dynamics to behavioral variation, Alzheimer’s disease pathology, cognitive performance, and age-related differences.^5,7,8^ Cognitive aging research therefore calls for representations that remain informative across populations and study designs while supporting assessment of current cognition, prediction of subsequent cognition, and characterization of within-person change.

Brain-imaging foundation models (FMs) aim to learn representations that can be reused across populations and research questions. Representation learning converts complex imaging data into numerical features through training, allowing patterns to be learned from the data rather than relying entirely on features specified in advance.^9^ Large-scale initial training, termed pretraining, enables features to be extracted from new scans without training a separate model from scratch for each study. Some brain-imaging FMs learn representations from FC matrices^10^, whereas others process regional rs-fMRI time series directly.^11^ Their pretraining objectives include reconstructing masked FC matrices or BOLD signals, predicting representations of withheld signals, or learning consistent features across different views of a recording.^10–15^ Model outputs can be summarized in different ways to obtain a representation for each scan, a step termed the readout. Researchers can extract these representations with model weights held fixed or further train the model for a particular population or task, a step termed post-training. These approaches may benefit cognitive aging research with limited samples, but their usefulness depends on whether the resulting representations retain information relevant to the intended measurement or prediction.

A recent benchmark evaluated fMRI FMs on clinical and demographic classification tasks, including Alzheimer’s disease classification on ADNI^16^ as well as age and sex classification in HCP-A,^17^ alongside decoding of cognitive task conditions and visual stimulus categories.^18^ Models showed inconsistent advantages over conventional FC for clinical classification, despite stronger performance on state-decoding tasks.^18^ These findings leave several questions unresolved for cognitive aging research: whether pretrained representations capture differences in memory and executive function, improve prediction of follow-up cognition beyond baseline clinical assessments, or characterize within-person changes accompanying an intervention. Addressing these questions also requires determining how readout choice affects downstream performance, given the sensitivity of brain–cognition associations to analysis choices.^19^

For use cases such as cognitive aging research, evaluating an FM requires distinguishing the information available in a pretrained model from how that information is extracted and used for downstream analysis. The encoder transforms rs-fMRI recordings into learned representations, while the readout aggregates these representations into scan-level embeddings. Comparing readouts with encoder weights held fixed allows us to assess how representation extraction affects concurrent cognitive associations and analyses of within-person change. These embeddings can then be compared with conventional FC features and representations from an FC-based FM, while their incremental value can be assessed by adding them to models based on clinical measures. Beyond readout choice, adaptation offers another way to tailor pretrained representations to aging populations. Post-training updates model weights using population- or task-specific data, whereas embedding alignment adjusts the distributions of extracted representations across cohorts without changing model weights. Comparisons with the corresponding frozen or unaligned representations can establish whether these approaches improve performance in the intended downstream setting.

In this study, we systematically evaluate nine configurations from five FM families that process regional rs-fMRI time series (**Table 1**) across two observational and five intervention aging cohorts (**Table 2** and **Fig. 1**), using checkpoints publicly available as of October 2026. We first benchmark frozen representations against conventional FC and a frozen FC-based FM for concurrent memory and executive-function associations (COG), age and sex prediction (DEMO), and diagnostic classification (DX) in observational cohorts, with baseline cognitive analyses also conducted in intervention cohorts. We then examine how readout choice affects cross-visit participant re-identification (Re-ID), using dimensionality matching, random-initialization controls, and regional signal-variability comparisons to interpret the information accessible through different readouts. We also evaluate readout effects on cognitive associations and assess cross-cohort cognitive transfer, distinguishing source-fitted predictors applied without refitting from locally fitted models. In intervention cohorts, we evaluate follow-up cognitive prediction from baseline measurements (COG-FU), cognitive-change associations (ΔCOG), and intervention-arm classification (ARM) using baseline and follow-up imaging. Clinical-only models provide references for assessing incremental imaging value in the longitudinal cognitive analyses. Finally, we evaluate post-training and embedding alignment against the corresponding frozen or unaligned representations under matched conditions. Together, these analyses provide a comprehensive assessment of the utility, generalizability, and adaptability of existing FMs for cognitive aging research, informing model selection and application across populations and cognitive outcomes.

**Table 1.**
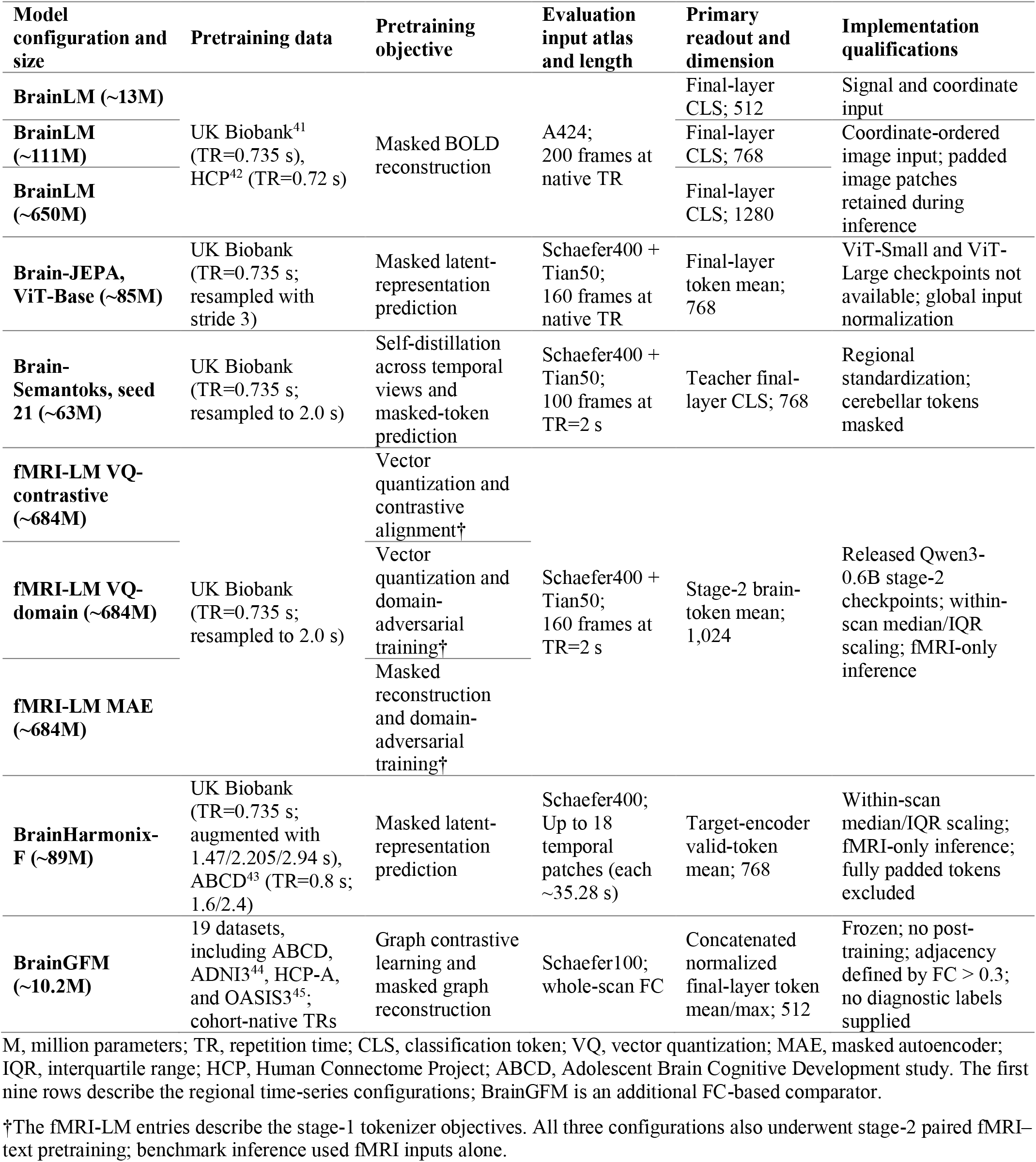
Model specifications and representation extraction.

| Model configuration and size | Pretraining data | Pretraining objective | Evaluation input atlas and length | Primary readout and dimension | Implementation qualifications |
|---|---|---|---|---|---|
| **BrainLM (~13M)** | UK Biobank<sup>41</sup> (TR=0.735 s), HCP<sup>42</sup> (TR=0.72 s) | Masked BOLD reconstruction | A424;<br>200 frames at native TR | Final-layer CLS; 512 | Signal and coordinate input |
| **BrainLM (~111M)** |  |  |  | Final-layer CLS; 768 | Coordinate-ordered image input; padded image patches retained during inference |
| **BrainLM (~650M)** |  |  |  | Final-layer CLS; 1280 |  |
| **Brain-JEPA, ViT-Base (~85M)** | UK Biobank (TR=0.735 s; resampled with stride 3) | Masked latent-representation prediction | Schaefer400 + Tian50;<br>160 frames at native TR | Final-layer token mean; 768 | ViT-Small and ViT-Large checkpoints not available; global input normalization |
| **Brain-Semantoks, seed 21 (~63M)** | UK Biobank (TR=0.735 s; resampled to 2.0 s) | Self-distillation across temporal views and masked-token prediction | Schaefer400 + Tian50;<br>100 frames at TR=2 s | Teacher final-layer CLS; 768 | Regional standardization; cerebellar tokens masked |
| **fMRI-LM VQ-contrastive (~684M)** | UK Biobank (TR=0.735 s; resampled to 2.0 s) | Vector quantization and contrastive alignment† | Schaefer400 + Tian50;<br>160 frames at TR=2 s | Stage-2 brain-token mean; 1,024 | Released Qwen3-0.6B stage-2 checkpoints; within-scan median/IQR scaling; fMRI-only inference |
| **fMRI-LM VQ-domain (~684M)** |  | Vector quantization and domain-adversarial training† |  |  |  |
| **fMRI-LM MAE (~684M)** |  | Masked reconstruction and domain-adversarial training† |  |  |  |
| **BrainHarmonix-F (~89M)** | UK Biobank (TR=0.735 s; augmented with 1.47/2.205/2.94 s), ABCD<sup>43</sup> (TR=0.8 s; 1.6/2.4) | Masked latent-representation prediction | Schaefer400;<br>Up to 18 temporal patches (each ~35.28 s) | Target-encoder valid-token mean; 768 | Within-scan median/IQR scaling; fMRI-only inference; fully padded tokens excluded |
| **BrainGFM (~10.2M)** | 19 datasets, including ABCD, ADNI3<sup>44</sup>, HCP-A, and OASIS3<sup>45</sup>; cohort-native TRs | Graph contrastive learning and masked graph reconstruction | Schaefer100; whole-scan FC | Concatenated normalized final-layer token mean/max; 512 | Frozen; no post-training; adjacency defined by FC > 0.3; no diagnostic labels supplied |

M, million parameters; TR, repetition time; CLS, classification token; VQ, vector quantization; MAE, masked autoencoder; IQR, interquartile range; HCP, Human Connectome Project; ABCD, Adolescent Brain Cognitive Development study. The first nine rows describe the regional time-series configurations; BrainGFM is an additional FC-based comparator.
†The fMRI-LM entries describe the stage-1 tokenizer objectives. All three configurations also underwent stage-2 paired fMRI–text pretraining; benchmark inference used fMRI inputs alone.

**Table 2.**
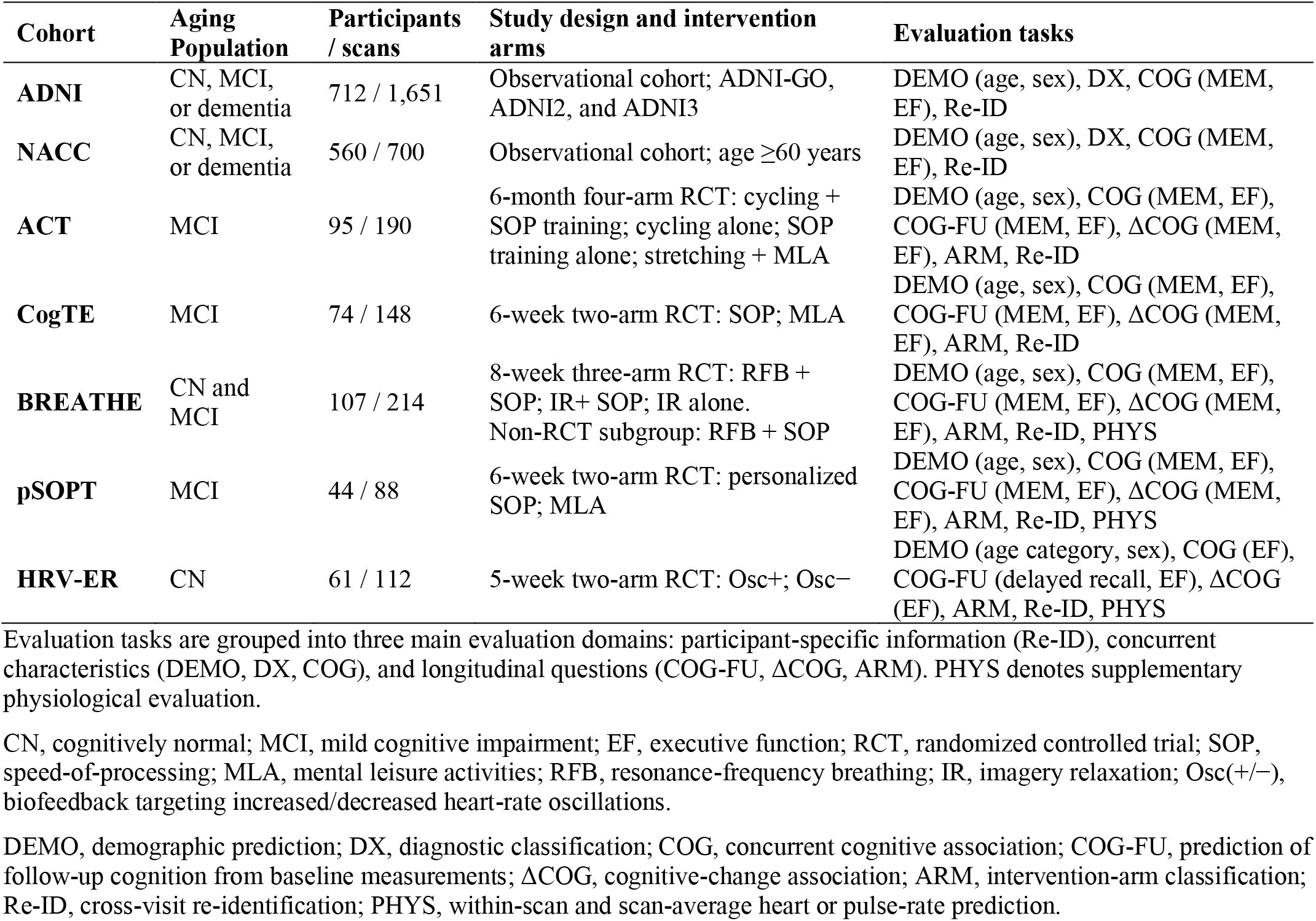
Cohorts and available evaluation tasks.

**Fig. 1.**
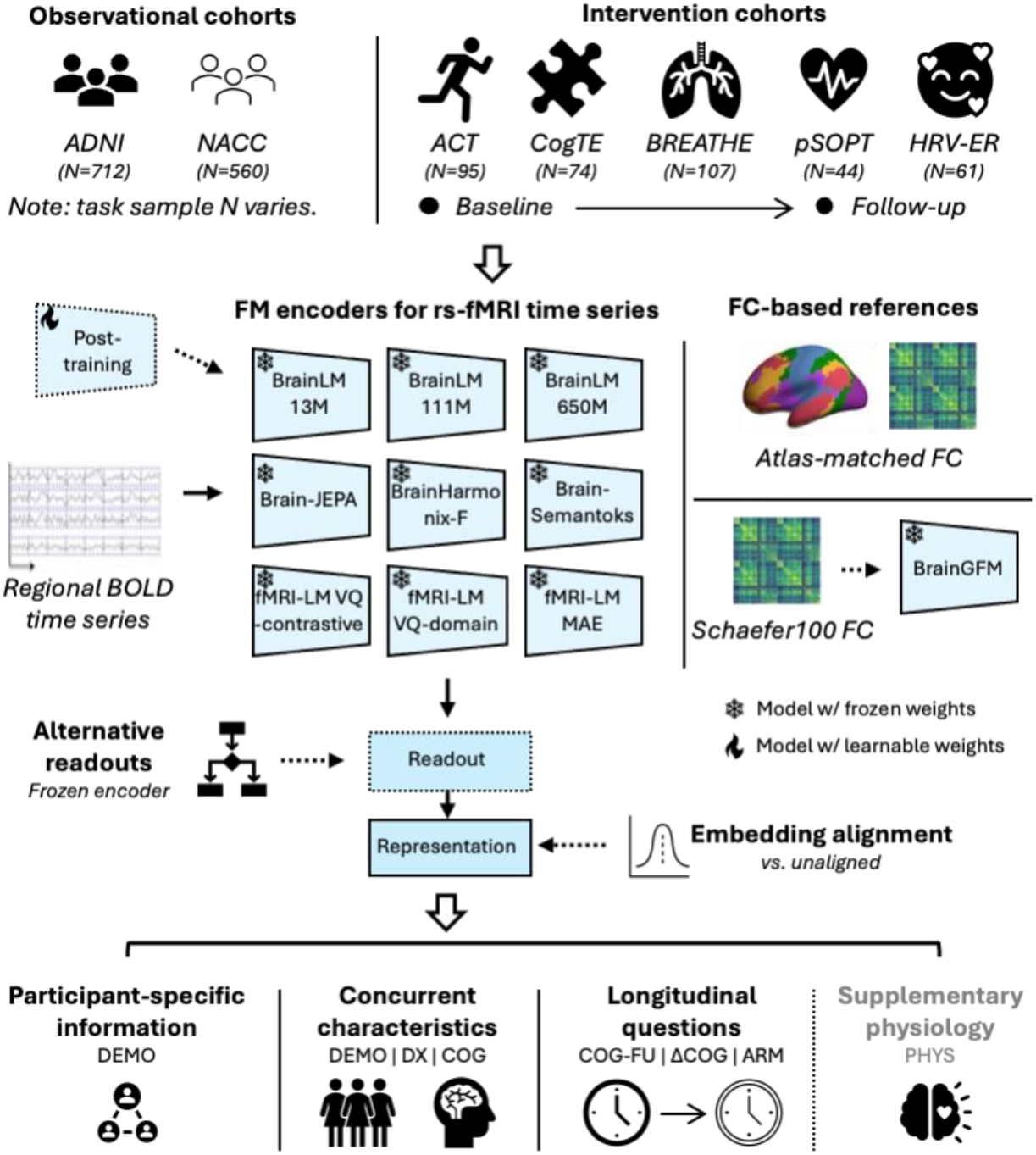
Study design and evaluation framework. Nine configurations from five foundation-model families that process regional rs-fMRI time series are evaluated across observational and intervention aging cohorts. Alternative readouts summarize encoder outputs with weights held fixed, post-training on ADNI or NACC updates encoders, and embedding alignment adjusts extracted representations. Conventional FC measures and frozen BrainGFM embeddings provide imaging references, with clinical or covariate references used in relevant diagnostic and cognitive analyses. Re-ID, cross-visit participant re-identification; DEMO, demographic prediction; DX, diagnostic classification; COG, concurrent cognitive associations; COG-FU, prediction of follow-up cognition from baseline measurements; ΔCOG, cognitive-change associations; ARM, assigned intervention-arm classification; PHYS, supplementary physiological evaluation.

## Methods

### Study design, cohorts, and outcomes

We evaluated FM representations derived from regional rs-fMRI time series across two observational aging cohorts, ADNI^16^ and NACC^20^, and five intervention aging cohorts, ACT,^3^ CogTE,^2^ BREATHE,^21^ pSOPT,^22^ and HRV-ER.^23^ **Table 2** summarizes their populations, study designs, interventions, and available evaluation tasks. ADNI included eligible scans from ADNI-GO, ADNI2, and ADNI3. NACC analyses required participants to be aged ≥60 years and the absolute interval between imaging and the associated cognitive assessment to be ≤6 months. BREATHE cognitive and intervention-arm analyses excluded the nonrandomized subgroup, and HRV-ER analyses used the older-adult subgroup.

ACT, CogTE, BREATHE, and pSOPT provided memory and executive-function (EF) assessments before and after intervention. HRV-ER provided pre- and post-intervention EF assessments, summarized as the mean of three NIH Toolbox standard scores, but lacked a baseline memory assessment. Its single Week-5 delayed-recall score was therefore evaluated as a subsequent memory outcome predicted from Week-2 MRI. HRV-ER also provided age categories rather than exact ages for demographic analyses and clinical covariates. Supplementary physiological analyses used electrocardiography (ECG)-derived heart rate in BREATHE and pSOPT and photoplethysmography (PPG)-derived pulse rate in HRV-ER. Detailed outcome definitions, eligibility criteria, and task-specific sample sizes are provided in **Supplementary Methods 1 and Table S1**.

### fMRI preprocessing and input construction

fMRI data were preprocessed using fMRIPrep,^24^ followed by a common regional time-series processing and extraction workflow with identical settings across cohorts. Processing included 6-mm full-width-at-half-maximum smoothing, detrending, 0.01–0.1-Hz filtering, and nuisance regression using available anatomical CompCor components, six motion parameters and their first derivatives, and cosine terms. Scans with excessive head motion were excluded during upstream quality control.

Regional signals were extracted without global signal regression using the A424, Schaefer400, or Schaefer400-plus-Tian parcellation required by each model (**Table 1**).^25–27^ No additional site harmonization was applied in the primary analyses, which evaluated pretrained representations under the acquisition heterogeneity present across cohorts. Embedding alignment was evaluated separately.

Model-specific input construction determined temporal coverage and scaling. BrainLM and Brain-JEPA retained native sampling intervals, whereas Brain-Semantoks and fMRI-LM inputs were resampled to a 2-s interval. Recordings were truncated or zero-padded to the required lengths. BrainHarmonix-F instead used temporal patches constructed at the native repetition time (TR). Zero-padded inputs were retained during inference unless excluded by model-specific masks. Brain-Semantoks used regional standardization and a pretrained missing-network embedding for unavailable cerebellar inputs. For fMRI-LM and BrainHarmonix-F, unavailable pretraining normalization statistics were replaced by within-scan regional median/interquartile-range scaling. Input lengths, normalization, and padding procedures are detailed in **Table 1, Supplementary Methods 2, and Table S2**.

### Foundation models and representations

We evaluated nine configurations from five pretrained FM families that accept regional BOLD time series: BrainLM 13M, 111M, and 650M; Brain-JEPA; Brain-Semantoks; fMRI-LM VQ-contrastive, VQ-domain, and MAE; and BrainHarmonix-F.^11– 15^ Models requiring volumetric or cortical-flat-map inputs were outside the scope of this benchmark.^18,28^ Model specifications, reported pretraining data, and representation dimensions are summarized in **Table 1**.

Primary scan-level representations used final-layer classification-token (CLS) embeddings for BrainLM and the Brain-Semantoks teacher encoder, mean-pooled regional and temporal tokens for Brain-JEPA, mean-pooled brain tokens for fMRI-LM, and mean-pooled valid tokens from the BrainHarmonix-F target encoder. fMRI-LM inference used brain inputs alone. Each configuration retained the same primary readout before and after post-training. During downstream representation extraction, model weights were fixed and random input masking was disabled. Model-specific masks for unavailable inputs remained active. Alternative readouts were evaluated separately using the same final encoder layer and fixed weights.

Reference representations included atlas-matched static FC and FC-Tucker,^29^ together with whole-scan Schaefer100 FC and frozen BrainGFM embeddings.^10^ Schaefer100 FC and BrainGFM used the same supplied FC matrices, and BrainGFM received no post-training. Additional input-window-matched FC controls assessed comparisons using the same observed temporal coverage as the corresponding FM. Checkpoint details, readout construction, and reference procedures are provided in **Supplementary Methods 3 and 6 and Tables S2–S4**.

### Downstream tasks and evaluation design

Methods used the same eligible samples and participant partitions within each comparison panel. Local prediction used ridge regression for continuous outcomes and logistic regression for categorical outcomes. Evaluation used five-fold cross-validation, keeping all scans from each participant in the same fold. In each iteration, one fold was held out for testing, and participants in the remaining four folds were divided approximately 80:20 for training and validation. This procedure was repeated with three random seeds. Outer folds were stratified by outcome classes or quintiles of continuous outcomes. Inner splits were randomized by participant, with additional arm stratification for paired-visit classification. Features were standardized to zero mean and unit variance, and principal component analysis (PCA) reduced dimensionality to limit prediction-model complexity. Feature transformations were fitted on the training partition and applied to validation and test participants. Hyperparameters were selected using validation performance, without using outer-test participants. An additional analysis of concurrent cognitive associations in ADNI and NACC jointly selected the final-layer readout and ridge regularization using inner-validation Spearman correlation (**Supplementary Methods 6; Table S4**).

Cross-cohort analyses distinguished transfer of an existing predictor from fitting a predictor locally. The cohort used to fit a transferred predictor is termed the source cohort, and the independent cohort in which it was evaluated is termed the target cohort. Source-fitted predictors were applied to target cohorts without refitting. Target-local analyses instead fitted downstream predictors using target-cohort training participants. Intervention leave-one-study-out (LOSO) evaluation trained predictors on the other cohorts and excluded held-out cohort outcomes from fitting and selection. HRV-ER delayed recall was evaluated separately by transferring memory predictors trained on the four other intervention cohorts (**Supplementary Methods 5; Table S6**).

Follow-up cognitive prediction (COG-FU) compared baseline imaging, clinical measures, and their combination. Paired-visit analyses evaluated cognitive-change associations (ΔCOG) and intervention-arm classification (ARM) using baseline representations, follow-up-minus-baseline differences, or their concatenation. Both visits used the same checkpoint and readout. For primary follow-up and paired-visit models, PCA was fitted to imaging features within each training partition, retaining at most five components to limit model complexity in the small intervention samples. Sensitivity analyses selected among caps of 5, 10, and 20 components using inner-validation performance. Clinical variables and task-specific procedures are detailed in **Supplementary Methods 4 and 10 and Tables S1, S7, and S8**.

Cross-visit Re-ID assessed whether each scan could be matched to another visit from the same participant. Accuracy was averaged across both retrieval directions, and within-versus between-participant similarity differences were evaluated separately (**Supplementary Methods 5; Table S5**). Supplementary physiological analyses assessed within-scan heart or pulse-rate variation and scan-average rates. Within-scan analyses centered features and rates by scan and adjusted for window position and visit; scan-average analyses adjusted for visit. All scans and windows from a participant remained in the same partition. Window representations extracted from full-context temporal tokens and those obtained by encoding each window independently were evaluated separately (**Supplementary Methods 9; Table S9**).

### Post-training and embedding alignment

Each configuration was post-trained separately on ADNI and NACC using supervised memory prediction, supervised EF prediction, and model-native self-supervised objectives, with three adaptation seeds. Here, the source cohort supplied the post-training data, and independent cohorts served as targets for external evaluation. Supervised post-training used low-rank adaptation (LoRA)^30^ (rank 8, α = 16) and mean squared-error loss to jointly optimize encoder adapters and a linear prediction head. Standardization and PCA were fitted to frozen training representations before adaptation and held fixed thereafter. PCA used a fixed cap of 100 components across configurations, further limited by training sample size and feature dimensionality. Downstream follow-up and paired-visit analyses used the separate PCA procedure described above, with a cap of five components.

Supervised checkpoints were selected by Spearman correlation on held-out validation participants from the source cohort, retaining the initial model if adaptation did not improve validation performance. Self-supervised post-training used model-native reconstruction, latent-prediction, or self-distillation objectives, with checkpoints selected by validation loss. For fMRI-LM, these objectives used imaging alone without repeating text-alignment pretraining. Evaluation within the source cohort fitted transformations, adapters, and prediction heads independently within each outer cross-validation fold. For external evaluation, encoder training and checkpoint selection used source-cohort data only. Results involving participants used for post-training were reported separately from independent-cohort evaluations (**Supplementary Methods 7; Table S10**).

Embedding alignment assessed whether adjusting representation distributions could improve cross-cohort performance without updating model weights. In source-fitted PCA coordinates, target embeddings were aligned to the source distribution by matching feature means and standard deviations or by matching covariance matrices using correlation alignment (CORAL).^31^ Alignment parameters were estimated from target calibration or training participants without their outcome labels and applied to held-out participants. Aligned and unaligned comparisons used the same underlying embeddings, evaluation samples, and participant partitions (**Supplementary Methods 8; Table S11**).

### Statistical, sensitivity, and exploratory analyses

Continuous outcomes were evaluated using Spearman correlation and prediction errors. Classification used binary or macro-averaged one-versus-rest ROC AUC, with balanced accuracy for HRV-ER age-category classification. Predictions were averaged across split seeds and, where applicable, adaptation seeds before scoring. Re-ID ensembles averaged similarity matrices before calculating accuracy.

Paired comparisons used 5,000 participant-level bootstrap resamples to estimate pointwise 95% confidence intervals and 20,000 participant-level swaps between methods for two-sided tests of performance differences. Swaps exchanged fixed predictions or participant-level Re-ID scores, keeping repeated observations and multiclass probability vectors together. These procedures did not refit models, and uncertainty estimates were therefore conditioned on fitted predictions and, for Re-ID, the fixed gallery. Absolute performance of primary frozen representations was assessed separately using 1,999 label permutations for within-cohort prediction, imaging-only follow-up cognitive prediction, and cognitive-change associations and intervention-arm classification using follow-up-minus-baseline representations. Each permutation regenerated outcome-stratified partitions and refitted the prediction pipeline. Repeated observations were permuted as participant blocks within visit-count strata. Re-ID used 20,000 random identity mappings, while within-scan physiological analyses used 1,999 scan-wise circular shifts with refitting. Absolute tests were one-sided (**Supplementary Methods 11 and 12; Tables S2 and S12**).

Holm correction was applied within defined comparison families. For label-permutation tests, families spanned configurations and eligible cohorts within each task. Above-chance evidence required a significant absolute test after correction in at least one cohort. Superiority over FC required a significant paired advantage over the designated FC reference after correction. Cross-cohort consistency required superiority to the same reference in at least two independent cohorts and positive differences in every evaluable cohort for the same task and protocol. Absolute significance and paired superiority were assessed separately, and nonsignificant differences were not interpreted as equivalence. Complete family definitions and analysis-specific inference procedures are provided in **Supplementary Methods 11 and 12 and Table S2**.

Sensitivity analyses examined final-layer readouts, dimensionality-matched fusion and Re-ID, available covariates, 80-frame inputs, unpadded scans, and participant sampling and weighting (**Supplementary Methods 6 and 10; Tables S4 and S7**). Embedding connectivity summarized pairwise similarities between final-layer spatial embeddings, averaged over temporal tokens where applicable and centered and normalized across feature dimensions. Across all nine configurations, Re-ID controls included five random initializations per configuration, regional BOLD standard deviations (SD), and input-window-matched FC. Primary and embedding-connectivity readouts were evaluated at native dimensionality and after projection to 128 dimensions. Temporal-order sensitivity used five shuffled versions of each scan, with the same temporal permutation applied across regions while preserving static FC, regional signal distributions, padding, and missing-input masks (**Supplementary Methods 14; Table S13**).

Supplementary SD comparisons extended to concurrent cognitive associations, follow-up cognitive prediction, cognitive-change associations, and intervention-arm classification. SD was calculated over model-matched observed input windows before model-specific regional normalization, excluding padding (**Supplementary Methods 15; Table S15**). Motion analyses assessed Re-ID after adjustment for mean framewise displacement (FD) across all nine configurations and seven cohorts, and separately evaluated prediction of scan-average FD from primary frozen embeddings using participant-grouped cross-validation (**Supplementary Methods 14 and 16; Table S13**).

## Results

### Concurrent information in frozen representations relative to FC references

Primary frozen representations showed above-chance performance in several demographic (DEMO), diagnostic (DX), and concurrent cognitive (COG) analyses, with different patterns across cohorts. All nine configurations exceeded chance for diagnostic classification and concurrent memory associations in NACC, whereas five configurations did so for each task in ADNI. For concurrent EF associations, five configurations exceeded chance in ADNI, but none did so in NACC (**Fig. S2A; Table S12**). **Fig. 2** summarizes performance relative to Schaefer100 FC across the seven cohorts, with cohort-specific profiles and absolute-test evidence provided in **Fig. S1 and Table S14**.

**Fig. 2.**
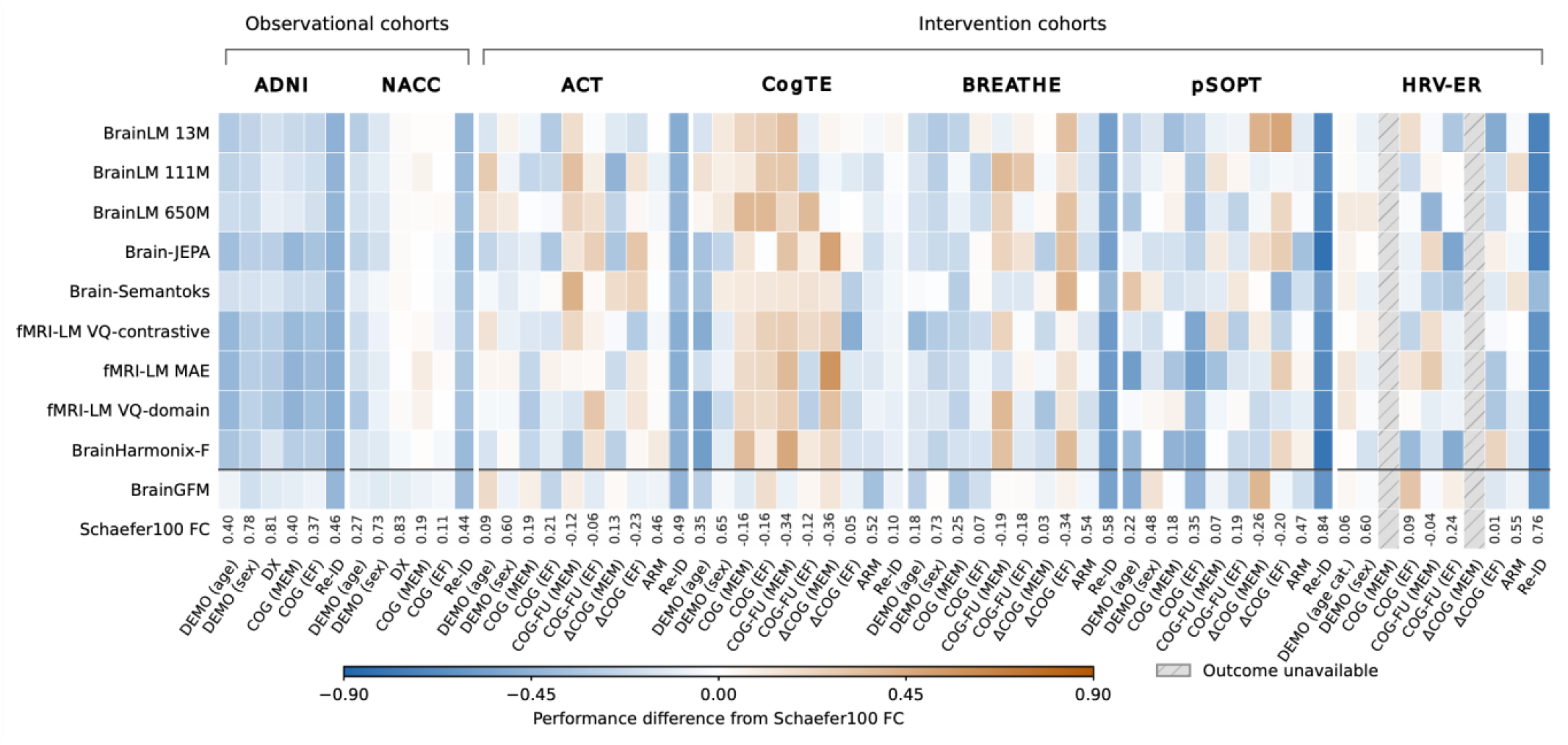
Performance of frozen representations across cognitive aging evaluation tasks relative to FC references. Primary readouts from nine time-series FM configurations and frozen BrainGFM representations are evaluated across two observational and five intervention cohorts. Cell color indicates the performance difference from Schaefer100 FC on matched samples, with warm colors indicating higher performance and cool colors indicating lower performance. Numerical values beneath each cohort report the corresponding Schaefer100 FC scores. ΔCOG and ARM used imaging-only follow-up-minus-baseline inputs. Metrics are Spearman correlation for continuous outcomes, ROC AUC for classification (macro-averaged one-versus-rest for multiclass arms), balanced accuracy for HRV-ER age-category classification, and bidirectional accuracy for Re-ID. Color differences should be interpreted within tasks because performance metrics differ across tasks. DEMO, demographic prediction; DX, diagnostic classification; COG, concurrent cognitive associations; COG-FU, prediction of follow-up cognition from baseline measurements; ΔCOG, cognitive-change association; ARM, intervention-arm classification; Re-ID, cross-visit participant re-identification; MEM denotes memory and EF denotes executive function.

Above-chance performance, however, did not generally translate into an advantage over FC. In ADNI, Schaefer100 FC outperformed all nine configurations for age prediction, sex and diagnostic classification, and concurrent memory associations after correction. Atlas-matched FC and FC-Tucker also outperformed all nine configurations for concurrent memory and EF associations in this cohort. In NACC, despite above-chance performance across all configurations for diagnosis and memory, none significantly outperformed Schaefer100 FC for either task after correction. The highest FM diagnostic AUC was 0.858, compared with 0.828 for Schaefer100 FC. A reference model incorporating demographic and acquisition variables achieved an AUC of 0.844, and adding FM representations did not significantly improve discrimination after correction. Across cohorts, no primary configuration met the criterion defined in Methods for consistent superiority over FC (**Tables S3 and S5**).

### Readout choice and participant-specific information

With primary frozen readouts, Brain-Semantoks exceeded chance in cross-visit Re-ID in six cohorts, whereas Schaefer100 FC exceeded chance in all seven. No primary FM readout significantly outperformed Schaefer100 FC (**Figs. S2B and S4; Tables S5 and S12**). Re-ID nevertheless improved when information from the same model was summarized differently. For BrainLM 13M, replacing final-layer CLS with embedding connectivity improved accuracy in all seven cohorts without updating model weights (all *P* = 0.012). These improvements persisted after projection to a common 128-dimensional space (**Fig. S3A and B**).

Random-initialization controls supported a contribution from pretraining to connectivity-based Re-ID. At native dimensionality, pretrained embedding connectivity achieved significantly higher accuracy than the corresponding randomly initialized representation in six of seven cohorts for BrainLM 13M, all seven for BrainLM 111M, and five for BrainLM 650M (**Fig. S3E**). In ADNI, BrainLM 13M achieved 31.1% accuracy with pretrained weights, compared with 0.6% averaged across five random initializations (*P* = 0.009). Its advantage over random initialization remained significant in six cohorts after dimensionality matching. Pretraining also increased the benefit of changing readouts: the accuracy gain from replacing CLS with embedding connectivity was greater with pretrained than randomly initialized weights in six cohorts (**Fig. S3C; Table S13**).

Whether alternative readouts outperformed conventional representations depended on the reference used. BrainLM 13M regional embedding norms outperformed input-window-matched A424 FC in ACT, CogTE, and BREATHE, reaching 87.7% accuracy in CogTE versus 65.1% for FC (*P* = 0.012). BrainLM 13M embedding connectivity also exceeded the matched FC reference in ADNI and BREATHE (**Table S4**). Regional BOLD SD, however, remained a strong identification reference. Across all nine configurations, embedding connectivity did not significantly outperform SD in any cohort at either native or matched dimensionality. In ADNI, SD achieved 45.3% accuracy, exceeding BrainLM 13M embedding connectivity (*P* = 0.016; **Table S13**). Temporal shuffling tested whether connectivity-based Re-ID was sensitive to input ordering despite the absence of a significant advantage over SD. Shuffling time points while preserving static FC and regional SD reduced BrainLM 13M accuracy from 66.3% to 39.1% in pSOPT and from 38.2% to 10.6% in HRV-ER (*P* = 0.017 and 0.006, respectively). Both reductions remained significant after dimensionality matching (**Fig. S3D**).

The readout advantage also persisted after adjustment for measured head motion. After adjustment for mean FD, BrainLM 13M embedding connectivity remained significantly more accurate than CLS in six of seven cohorts across all three calibration splits, at both native and matched dimensionality. Effects in other configurations varied across cohorts and splits, and no adjusted connectivity readout significantly outperformed its adjusted input-window FC reference (**Fig. S3F; Table S13**). Separately, primary embeddings predicted scan-average FD, with correlations ranging from 0.370 to 0.534 across the three BrainLM configurations in ADNI and from 0.621 to 0.701 across the three fMRI-LM configurations in HRV-ER (**Table S13**).

### Cognitive associations and longitudinal utility across cohorts

Readout changes also improved concurrent cognitive associations, although the gains varied across tasks and cohorts. In ADNI, flattening BrainLM 13M final-layer regional embeddings increased the correlation between predicted and observed memory scores from 0.185 with CLS to 0.390, and the corresponding EF correlation from 0.133 to 0.327 (both *P* = 0.003; **Fig. 3A and B**). Unlike the Re-ID comparisons, selected cognitive analyses showed advantages over regional BOLD SD. In the matched NACC memory analysis, fMRI-LM MAE embedding connectivity achieved ρ = 0.352, compared with 0.188 for window-matched SD (*P* = 0.020; **Table S15**). Advantages over FC were more limited. An alternative fMRI-LM MAE readout outperformed Schaefer100 FC for NACC memory, but no alternative readout significantly exceeded atlas-matched FC, FC-Tucker, or input-window-matched FC in the observational cognitive panel (**Table S4**).

**Fig. 3.**
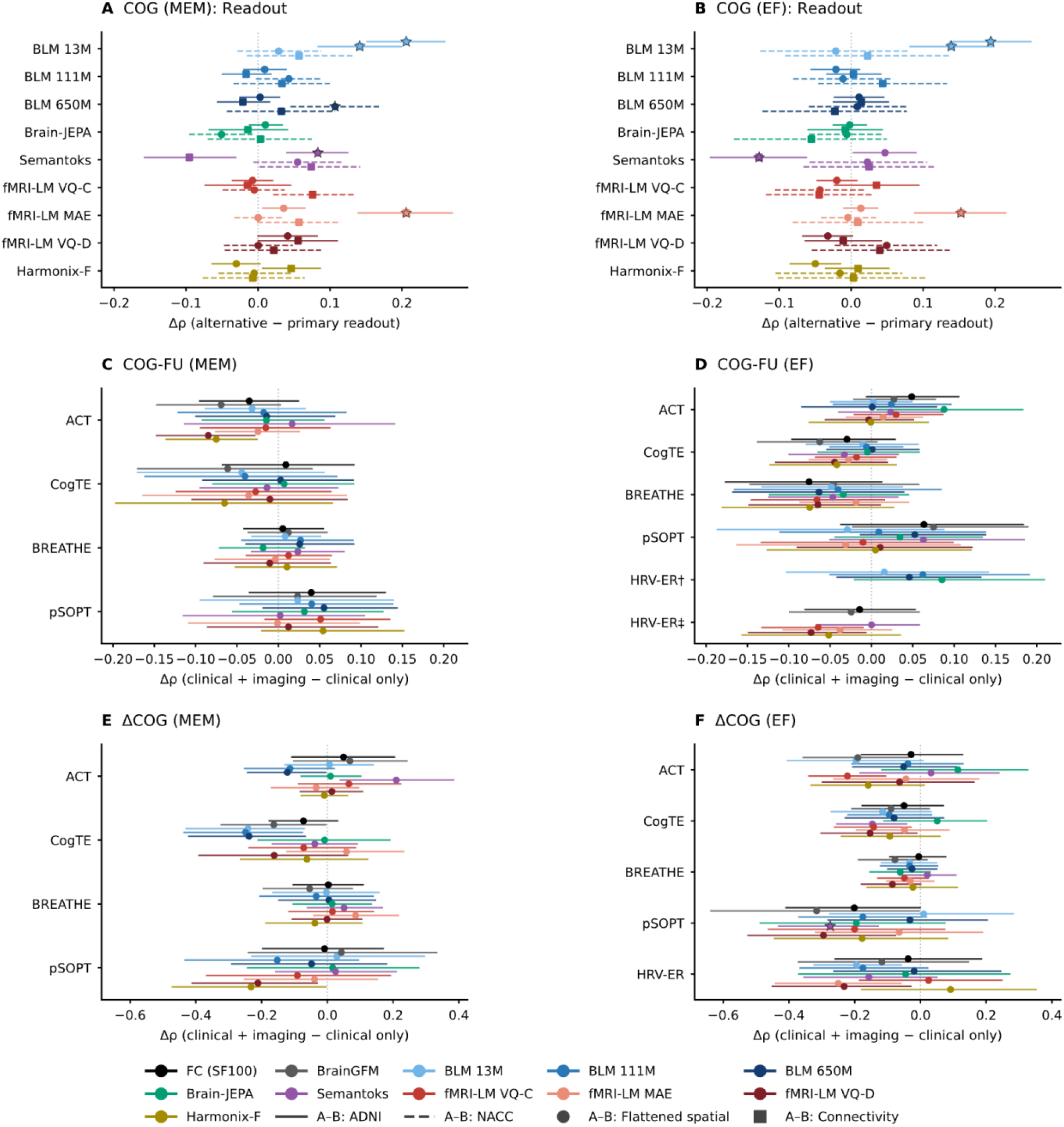
Readout effects on concurrent cognitive associations and incremental imaging value for longitudinal cognition. **(A and B)** Differences in memory and executive-function (EF) association performance between alternative final-layer readouts and each model’s primary readout in ADNI and NACC, with encoder parameters held fixed. Positive values favor the alternative readout. **(C and D)** Differences in follow-up cognitive prediction between models combining baseline clinical and imaging measurements and clinical-only models. **(E and F)** Differences in cognitive-change association performance between models combining clinical measures and follow-up-minus-baseline imaging representations and clinical-only models. Positive values in **C–F** indicate incremental imaging value. In HRV-ER, clinical-only correlations were 0.730 (†) and 0.818 (‡). Both analyses used baseline EF, age category, sex, and intervention arm in the same 55 participants and identical cross-validation splits but differed in how categorical variables were encoded (**Supplementary Methods 4**). Intervals are pointwise conditional 95% participant-bootstrap confidence intervals; stars indicate Holm-adjusted *P* < 0.05, including both increases and decreases.

These within-cohort gains did not translate uniformly to cross-cohort transfer. Changing the readout improved selected source-to-target transfers but reduced performance in others (**Tables S4 and S6**). Primary-readout transfer results are shown in **Fig. S5A and B**. In intervention-cohort LOSO evaluation, where predictors were trained on the other cohorts, neither the primary frozen FMs nor BrainGFM significantly outperformed Schaefer100 FC after correction. The same held for the separate transfer of memory predictors to HRV-ER delayed recall (**Fig. S5C and D; Table S6**).

Longitudinal analyses assessed both follow-up cognitive prediction from baseline measurements and associations involving changes between visits. No primary frozen configuration exceeded chance after correction for imaging-only follow-up cognitive prediction, or for cognitive-change associations and intervention-arm classification using follow-up-minus-baseline representations (**Fig. S2C; Table S12**). Some relative advantages nevertheless emerged. Brain-Semantoks outperformed Schaefer100 FC for ACT follow-up memory prediction, although its performance did not exceed chance in the corrected absolute test. In HRV-ER, longitudinal representations improved intervention-arm classification relative to baseline-only imaging but did not exceed chance or outperform Schaefer100 FC after correction (**Fig. S6A; Tables S8 and S12**).

Clinical-only models predicted follow-up cognition with correlations of 0.571–0.778 for memory and 0.583–0.818 for EF across eligible cohorts. Adding baseline imaging did not significantly improve follow-up prediction, and adding longitudinal imaging did not significantly improve cognitive-change associations beyond clinical measures after correction in the primary matched analyses (**Fig. 3C–F; Tables S8 and S10**). Complete baseline, change, and combined-input comparisons are reported in **Fig. S7 and Table S8**, with alternative-readout intervention analyses and PCA sensitivity results in **Tables S4 and S7**, respectively.

### Post-training and embedding alignment

Holding the primary readout fixed, we examined whether updating model weights improved cognitive performance. Supervised post-training improved memory associations in two within-source comparisons under participant-level cross-validation (**Fig. 4A and B**) and in two independent-cohort settings. The external gains occurred with both source-fitted transfer and target-local fitting. ADNI EF post-training improved Brain-Semantoks memory prediction in NACC when the ADNI-fitted predictor was transferred without refitting (Δρ = 0.304, 95% CI, 0.204–0.405, *P* = 0.009; **Fig. 4C**). NACC memory post-training also improved fMRI-LM VQ-contrastive baseline memory associations in BREATHE, where the downstream predictor was fitted locally (**Fig. 4D**). No improvement in follow-up cognitive prediction, cognitive-change associations, or intervention-arm classification was significant after correction (**Tables S8 and S10**). Complete primary-readout ADNI–NACC transfer comparisons are shown in **Fig. S8A and B**.

**Fig. 4.**
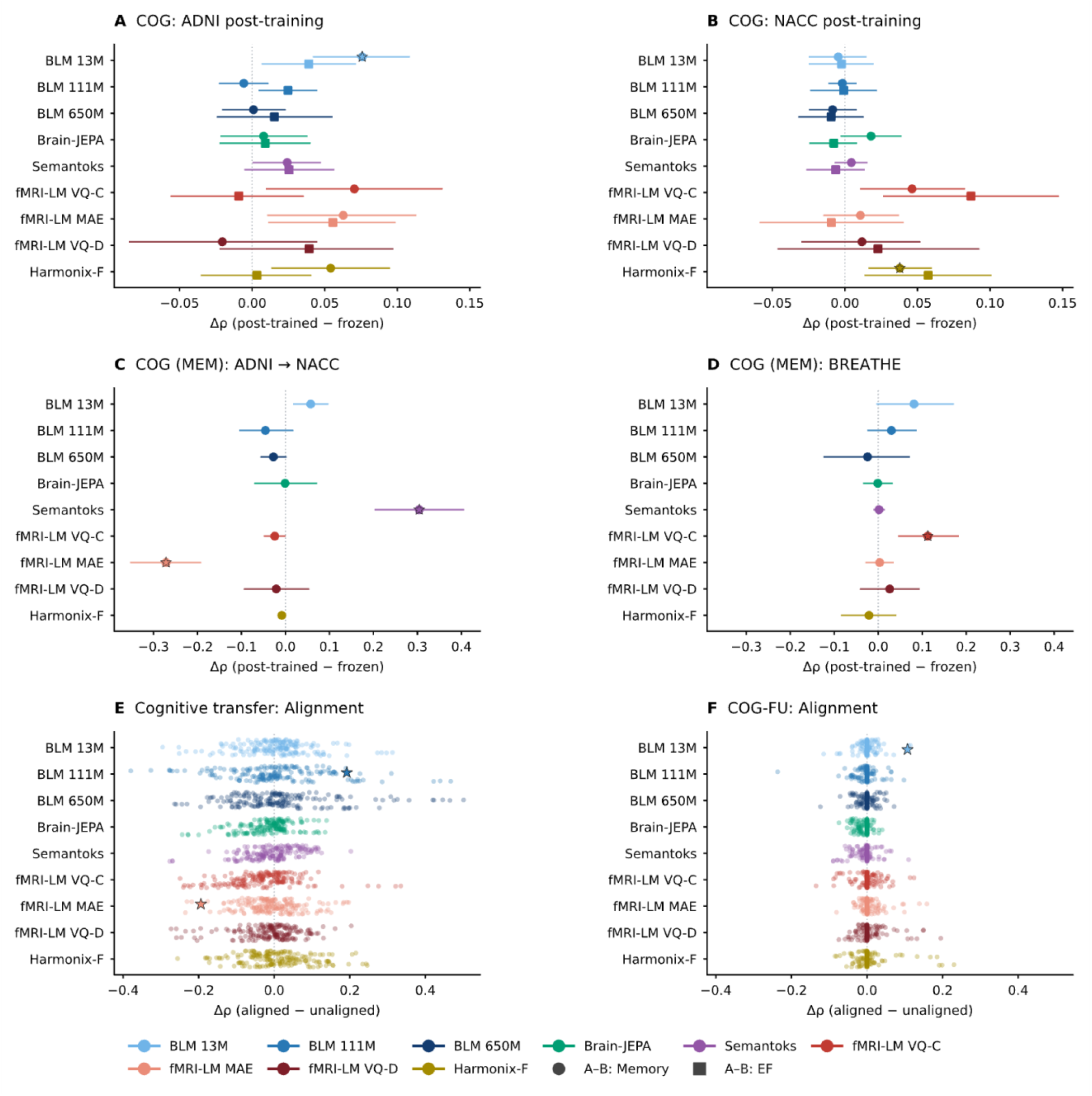
Effects of post-training and embedding alignment on cognitive evaluation performance. **(A and B)** Supervised post-training compared with the corresponding frozen encoders for concurrent memory and executive-function (EF) associations in ADNI and NACC, evaluated using fold-specific encoders. **(C)** ADNI-to-NACC memory evaluation after ADNI EF post-training, with the source-fitted statistical model applied without refitting. **(D)** Concurrent baseline-memory evaluation in BREATHE after NACC memory post-training, with the statistical model fitted locally in BREATHE. **(E and F)** Aligned-versus-unaligned performance differences across all 3,168 evaluated mean/standard-deviation and covariance-alignment comparisons, covering source-fitted model transfer and locally fitted follow-up cognition or delayed-recall analyses, respectively. Each point represents a comparison. Positive differences favor post-training or alignment. **A–D** show paired differences and pointwise conditional 95% confidence intervals. Stars throughout indicate Holm-adjusted *P* < 0.05 for either an increase or a decrease.

A separate exploratory panel examined whether changing the readout remained beneficial after post-training. This panel used locally fitted predictors in independent target cohorts and compared post-trained and frozen representations using the same readout. None of these post-trained-versus-frozen differences was significant after correction within this panel (**Fig. S8C; Table S10**). However, replacing the post-trained models’ primary readouts with embedding connectivity improved selected cognitive associations (**Fig. S8D**). These comparisons attributed the observed gains within this panel to changing the readout rather than demonstrating a benefit from post-training.

Embedding alignment provided a separate test of whether adjusting extracted representations could improve performance without updating model weights. Improvements survived correction in one fixed-source memory-transfer setting and one locally fitted follow-up EF setting (**Fig. 4E and F**). In paired-visit analyses, alignment also improved one intervention-arm comparison relative to its matched unaligned control, but not relative to the original primary pipeline or Schaefer100 FC (**Fig. S9; Table S11**). Additional input-length and physiological results are provided in **Fig. S10 and Table S7**, and **Fig. S11 and Table S9**, respectively.

## Discussion

Foundation models have the potential to generate brain representations that can be reused in individual studies.^32,33^ Across seven aging cohorts, pretrained rs-fMRI representations supported cross-visit re-identification and selected concurrent cognitive associations, but their utility depended on how model outputs were summarized. Changing the readout improved performance without updating model weights, showing that evaluation through a single primary readout can provide an incomplete assessment of a pretrained model. Readout improvements did not consistently establish superiority over FC, and the primary longitudinal analyses did not demonstrate added value beyond baseline clinical measures (**Fig. 3; Tables S4 and S8**). For cognitive aging research, these findings distinguish evidence that a representation contains useful information from evidence that it improves on established measures for the intended task. This emphasis on demonstrated utility is consistent with broader recommendations for evaluating biomedical foundation models.^34^

Readout effects were particularly consistent for BrainLM 13M Re-ID. Replacing final-layer CLS with embedding connectivity improved Re-ID across all seven cohorts, including after dimensionality matching (**Fig. S3A and B**). Random-initialization controls supported a contribution from pretraining to this gain and to identification performance in several other configurations (**Fig. S3C and E**). These findings show that readout choice affects access to individual information, but they do not establish that this information exceeds what simpler signal summaries provide. Across all nine configurations, embedding connectivity did not significantly outperform regional BOLD SD for Re-ID (**Table S13**). Temporal shuffling nevertheless reduced Re-ID in selected settings while preserving static FC and regional SD (**Fig. S3D**). This sensitivity to input ordering does not isolate meaningful neural dynamics, because shuffling also changes the distribution of inputs presented to the model. The relevance of these representations to cognition therefore requires separate evaluation. In cognitive analyses, selected FM representations outperformed SD, whereas others performed worse (**Table S15**). Strong Re-ID performance alone is consequently insufficient to guide representation choice for cognitive aging research, so readouts should be evaluated against the intended cognitive outcome.

Performance in observational studies does not establish whether a representation is useful for intervention research. In NACC, diagnostic discrimination did not translate into significant improvement beyond measured demographic and acquisition covariates after correction. Distinguishing individuals or diagnostic groups also does not establish sensitivity to change within a person. In intervention cohorts, baseline clinical assessments strongly predicted follow-up cognition, and the primary analyses did not demonstrate additional predictive value from imaging. Longitudinal representations improved selected intervention-arm comparisons relative to baseline imaging, but cognitive-change associations and intervention-arm classification using primary frozen change representations did not exceed chance after correction. Their usefulness for measuring intervention-associated change therefore remains unresolved. Because these analyses incorporated follow-up imaging, they evaluated longitudinal associations rather than pretreatment prediction of individual benefit. Establishing intervention relevance will require distinguishing treatment-related changes from measurement variability and changes that would have occurred without treatment. It also requires separating representation development from trial evaluation and specifying how intervention outcomes inform model selection and interpretation.^9^

Model selection should begin with the recordings and outcomes available in the intended study. Equal frame counts can span different durations across repetition times, and truncation or padding changes the observed information entering a model. Although longer recordings can improve connectome-based prediction,^35^ our input-length findings provided no general basis for preferring longer or shorter FM inputs. Differences in cognitive assessments and their reliability also constrain comparisons across studies.^36^ Within these constraints, readout choice provides an opportunity to improve performance without updating model weights. Selecting readouts within training partitions improved selected cognitive predictions over primary readouts in held-out ADNI participants, supporting the evaluation of readout choice as part of local model selection (**Table S4**). When adaptation is considered, its benefit should be assessed against a suitable frozen representation. Post-training improved selected source-cohort and independent-cohort predictions, but gains did not extend consistently across tasks, in line with evidence that better performance in an adaptation setting need not imply better generalization.^37^ Embedding alignment also yielded improvements in selected settings without updating model weights. For smaller studies, evaluating these options requires keeping evaluation participants separate from those used for fitting and selection, while retaining enough data to assess whether any benefit is reliable.

Several limitations affect these conclusions. Our findings are specific to the evaluated checkpoints and procedures, and modest intervention samples limited the precision of performance estimates.^38^ Comparisons also differed in the amount of observed data available to each approach, as whole-scan Schaefer100 FC and the restricted inputs used by several FMs were not directly equivalent. Although atlas- and window-matched FC analyses helped address this imbalance, input equivalence could not be achieved across all comparison panels. In addition, unavailable pretraining normalization statistics necessitated alternative scaling for some models. Motion regression and covariate adjustment may not have fully accounted for residual motion or acquisition-related variation, and site effects were not explicitly harmonized. Finally, because population and acquisition characteristics varied together, differences in transfer performance could not be attributed to any single source of mismatch.

Overall, these findings identify priorities for developing and evaluating brain-imaging FMs for cognitive aging research. Broader representation of older adults across cognitive and clinical states, together with longitudinal observations, could provide a basis for learning features relevant to cognitive aging. Future models should also incorporate training objectives and interpretable readouts that preserve and capture meaningful within-person changes over time. Equally important, independent evaluation across cohorts and analytical pipelines is needed to determine whether these improvements generalize beyond the settings in which the models were developed. This emphasis on rigorous and standardized validation aligns with NIH’s PRIMED-AI program, which includes a dedicated Validation Center to support independent assessment.^39^ Ultimately, the value of brain-imaging FMs for cognitive aging research should be determined not only by their general predictive performance, but also by their ability to support reliable measurement and prediction of aging-related changes relevant to specific scientific questions.^40^

## Data availability

ADNI data are available through the LONI Image and Data Archive, and NACC data are available through the NACC data request process, subject to their respective access approvals and data use agreements. HRV-ER data are publicly available through OpenNeuro, dataset ds003823, version 1.3.5. Access to ACT, CogTE, BREATHE, and pSOPT data may be requested from the [StanfordCogTlab](https://cogtlab.stanford.edu/), subject to applicable institutional approvals and data use agreements. Aggregate results and numerical data underlying the figures are provided in **Tables S1–S15 and Datasets S1–S16** and accompanying supplementary files.

## Code availability

Custom code for representation extraction, post-training, downstream evaluation, statistical testing, and figure generation is available in our [GitHubrepository](https://github.com/ymliu17/brainfm). Original pretrained checkpoints were obtained from the model developers. Checkpoint specifications and implementation details are provided in the **Supplementary Methods 3 and 7** and **Tables S2 and S10**. Requests for post-trained adapters should be directed to the corresponding author and are subject to applicable data-use conditions.

## Author contributions

Y.M.L. conducted the analyses and drafted the manuscript. M.A. performed data preprocessing. K.Z. contributed to the design and refinement of the experimental protocols. Y.Z. and F.V.-L. supervised the study, guided its conceptualization and methodology, and contributed to interpretation of the findings. All authors critically reviewed and revised the manuscript and approved the final version.

## Acknowledgments

This work was supported by the National Institutes of Health (R13AG099775, P30AG102665, R33AG073356, P30AG066515, R61AG081723, R01NR015452, R21AG080425, and R01MH129694). Additional support was provided by the Stanford Knight Initiative for Brain Resilience, Stanford Institute for Human-centered AI, the Rosenkranz Foundation (grant RAR1-03), and the Eagles Autism Foundation.

## Competing interests

The authors declare no competing interests.

## Notes

### Competing Interest Statement

The authors have declared no competing interest.

