## Supporting information for "Benchmarking foundation models for resting-state functional magnetic resonance imaging time series in cognitive aging research"

SI Appendix: supplementary methods, results, references, Table S14, and Figs. S1–S11. Numerical Tables S1–S13 and S15 and supporting records are supplied in Datasets S1–S16.

### Contents

Supplementary Methods 1. Cohorts, outcomes, and analytical samples

Supplementary Methods 2. Preprocessing, temporal coverage, and model-specific inputs

Supplementary Methods 3. Foundation-model configurations and reference representations

Supplementary Methods 4. Local prediction, clinical references, and paired-visit outcomes

Supplementary Methods 5. Cross-cohort transfer and cross-visit re-identification

Supplementary Methods 6. Final-layer readouts and the focused adaptation interaction

Supplementary Methods 7. Supervised and model-native self-supervised post-training

Supplementary Methods 8. Target-to-source embedding alignment

Supplementary Methods 9. Physiological outcomes and window representations

Supplementary Methods 10. Fusion, input context, covariates, and PCA sensitivity

Supplementary Methods 11. Statistical inference, multiplicity, and consistency

Supplementary Methods 12. Absolute tests and evidence matrix

Supplementary Methods 13. Data, code, and supporting materials

Supplementary Methods 14. Additional Re-ID controls

Supplementary Methods 15. Regional signal variability beyond Re-ID

Supplementary Methods 16. Prediction of scan-average framewise displacement

Supplementary Results; supplementary table descriptions and Table S14; Figs. S1–S11; supplementary references.

### Supplementary Methods 1. Cohorts, outcomes, and analytical samples

Task labels are COG, concurrent cognitive associations; COG-FU, follow-up cognitive prediction; ΔCOG, cognitive-change associations; ARM, intervention-arm classification; DEMO, age and sex prediction; DX, diagnostic classification; Re-ID, cross-visit re-identification; and PHYS, physiological prediction. This appendix describes the evaluation of nine configurations from five resting-state fMRI foundation-model families across two observational and five intervention cohorts. Numerical Tables S1–S13 are supplied in Datasets S1–S13, respectively. Dataset S14 contains detailed PCA-selection records accompanying Table S7; Dataset S15 contains source data for main Figs. 2–4 and Figs. S1–S11. Each workbook includes a contents sheet and field definitions. Analysis-specific sample counts accompany the numerical comparisons. Blank numerical cells denote unavailable or inapplicable estimates, not zero. The cohort-specific evidence matrix is provided in Table S14, with numerical support in Dataset S12. Table S15 contains the regional-SD prediction controls and is supplied in Dataset S16.

ADNI contributed scan-associated memory and executive-function (EF) composites, age, sex, and diagnostic labels from records identified as ADNI-GO, ADNI2, or ADNI3. Repeated scans remained grouped by participant. The primary diagnosis distinguished normal cognition from dementia; intermediate diagnostic states were excluded from that binary task. These scan-visit labels should not be interpreted as independently verified etiologic diagnoses. Primary cognitive panels contained 1,651 memory observations from 712 participants and 1,643 EF observations from 710 participants. Task-specific counts are reported rather than a single cohort denominator (1).

NACC cross-sectional analyses selected one scan per participant, required age of at least 60 years, and retained an absolute imaging–cognitive-assessment interval of at most six months. Memory and EF used the supplied cognitive composites. Primary diagnosis coded NACCUDSD=1 as normal cognition and NACCUDSD=4 as all-cause dementia. The separate AD-etiology task required concordant NACCUDSD, NORMCOG, and DEMENTED fields at the associated clinical month, with NACCALZD=1 for dementia cases. Other diagnostic states were excluded. Memory, EF, and primary diagnosis samples comprised 544, 477, and 414 participants; the AD-etiology sample contained 196. A separately assembled repeated-scan NACC sample supported identification and self-supervised training (2).

ACT, CogTE, BREATHE, and pSOPT supplied repeated memory and EF measures. Scores were used in their supplied cohort-specific units; identical endpoint names do not imply identical instruments across cohorts. ACT follow-up corresponded to the six-month visit and CogTE to the six-week visit. BREATHE cognitive and arm analyses excluded the nonrandomized cognitively unimpaired subgroup. Its identification and physiological samples were assembled separately and could include that subgroup. The Schaefer100 archive mapped ACT T3_6mo to benchmark T2 and used BREATHE run-01 (3–6).

The recorded label builders used delayed-recall measures for memory rather than requiring a multi-test memory composite in every intervention cohort. BREATHE used BVMT-R delayed-recall T scores from sessions 1 and 2; both were expressed on the session-1 mean/SD scale, with the supplied EF pre/post composites retained separately. pSOPT used BVMT-R delayed recall and the supplied EF composite. Its delayed-recall boundary codes <20 and >80 were decoded to 19 and 81 in the existing label preparation. These are coded censored observations, not missing outcomes imputed by a downstream model. The same decoded scores underlie the reported continuous change magnitudes; uncertainty due to censoring was not separately modeled. Table S1 records the exact source column mappings.

HRV-ER used the older-adult subgroup of OpenNeuro ds003823, version 1.3.5. MRI visits occurred in Weeks 2 and 7. The study-defined EF composite was the arithmetic mean of the uncorrected standard scores for Flanker, List Sorting Working Memory, and Pattern Comparison Processing Speed, requiring all three at each visit. It is not an official NIH Toolbox EF composite. The NIH Toolbox computed-score fields were not relabeled as domain composites. A single Week-5 delayed picture-recall assessment supplied a distinct memory endpoint: the number of distinct correctly recalled images across valences (possible range 0–36) after Week-4 encoding. Duplicate correctly coded images were counted once; explicit zero recall was retained and missing tasks were not assigned zero. Week-2 MRI supported prospective prediction, whereas Week-7 MRI supported a retrospective association. Baseline episodic-memory assessments were unavailable, so neither memory change nor a baseline MEM covariate was constructed. Exact age was not released; available age categories were retained as categorical variables (7).

Common-sample comparisons used the intersection of available model inputs and task-specific finite outcomes, with the same participant partitions for all methods within a panel. The common Schaefer100 panel excluded one ambiguous CogTE baseline export and one pSOPT follow-up scan without a confirmed archive match. Cognitive eligibility was not imposed on the separate physiological panel. Sparse constant parcels were retained under the specified signal-availability policy rather than removed solely because at least one parcel was constant. The input audit rejected nonfinite recordings and extensive constant-region coverage (more than 5% of parcels, using a standard-deviation threshold of 10^-6); the strict no-constant-parcel flag was descriptive. This numerical check did not establish spatial registration or motion quality. Model-input ROI ordering was checked against the retained atlas mappings.

### Supplementary Methods 2. Preprocessing, temporal coverage, and model-specific inputs

fMRI data were preprocessed using fMRIPrep, followed by a common regional time-series processing and extraction workflow with identical settings across cohorts. Scans with excessive head motion were excluded during upstream quality control. Processing included 6-mm full-width-at-half-maximum spatial smoothing, detrending, 0.01–0.1-Hz filtering, and nuisance regression using available anatomical CompCor components, six rigid-body motion parameters and their first derivatives, and cosine drift terms. Regional signals were extracted without global signal regression. No additional site harmonization was applied in the primary analyses, which evaluated representations under the acquisition heterogeneity present across cohorts. Embedding alignment was evaluated separately as described in Supplementary Methods 8. The benchmark reused the resulting regional signals. Acquisition information is provided in Table S1 (8).

BrainLM received A424 signals (424 regions); Brain-JEPA received Schaefer400 plus Tian-S3 signals (450 regions). Input construction first retained real samples, then cropped the first 200 frames for BrainLM or 160 for Brain-JEPA. Each region was centered over its retained real frames before right zero-padding a shorter recording to the required length. Storage padding was removed using recorded valid lengths. No additional cohort-dependent amplitude correction, per-region variance normalization, temporal stretching, or TR resampling was applied by this common construction step. Native TR was retained, so frame-matched inputs were not necessarily duration-matched across cohorts (9–11).

BrainLM 13M used the native signal-vector and coordinate input path. For the 111M and 650M releases, regional rows were ordered by their Y coordinates, divided by the released scaling constant 5.6430855, repeated into three channels, and padded symmetrically with -1 to the model image dimensions (432×432 and 434×434, respectively). This image padding was distinct from the preceding temporal zero-padding. Primary inference retained all patches, including fully image-padded patches, and used deterministic token order. An all-zero inference mask disabled token removal; this differs from excluding architectural padding from the reconstruction loss. Brain-JEPA used its official global normalization, with one mean and standard deviation across the full regional-time input, and the released 450-region gradient-position table. These model-specific transformations were retained before and after adaptation.

Brain-Semantoks and the three fMRI-LM variants used polyphase resampling to TR=2 s. The released ROI/network order was reconstructed from the Schaefer atlas correspondence. fMRI-LM placed the 50 subcortical regions before the 400 cortical regions. Brain-Semantoks used its cortical/subcortical network ordering; the unavailable cerebellar network was represented by the pretrained missing-network embedding.

For Brain-Semantoks, regional signals were standardized within scans; for fMRI-LM, each region was centered by its scan median and divided by its interquartile range (floor 10^-6). Constant regions became zero. Normalization used the retained up-to-180 resampled frames, followed by deterministic first-100-frame Brain-Semantoks or first-160-frame fMRI-LM inference. Short arrays were right-padded with zeros. These models did not receive a temporal attention mask that excluded every padded sample. The missing-network mask for Brain-Semantoks is a different operation.

BrainHarmonix-F used the 400 cortical Schaefer regions at native TR. Patch length was round(35.28/TR) frames, with a maximum of 18 patches. Regional median/IQR normalization used the retained real recording. Fully padded tokens were excluded from the encoder/readout; a partially filled terminal patch could contain zeros. Fixed sinusoidal positions were regenerated for the supported 18-patch length. The released target encoder weights were otherwise restored strictly.

The pretraining-corpus normalization statistics required for exact fMRI-LM and BrainHarmonix-F input replication were unavailable. Per-scan median/IQR normalization was therefore a benchmark implementation choice. These results evaluate the documented implementations, not an exact reproduction of every pretraining transformation. Temporal zero-padding remained visible to encoders except where the implemented validity masks excluded whole tokens; it was not assumed to be uniformly masked across architectures.

The Schaefer100 comparison used supplied whole-scan Pearson FC matrices without GSR. Direct FC and BrainGFM shared exactly the same matrix for each scan, but these matrices were not matched to the 200-/160-frame model contexts. Their atlas and recording coverage therefore differed from the time-series model inputs. FC-Tucker could not be calculated for Schaefer100 because regional time series were not supplied; second-order FC matrices do not recover the required co-skewness information.

Atlas-matched A424 FC/FC-Tucker used the first 200 native frames; Schaefer450 references used 160 native frames, and the BrainHarmonix-F atlas reference used the 400 cortical regions from that same 160-frame input. Separate input-window controls reconstructed FC from each model's documented retained context, including resampling or patch duration where applicable. These are distinct reference panels. Whole-scan Schaefer100 FC, atlas-matched fixed-frame FC, and model-input-window FC should not be interchanged when quoting a score or P value. Available native TRs were 3 s for ADNI, 0.607 or 3 s for NACC, 1.03 s for ACT, 2.5 s for CogTE, 1.01 s for BREATHE, 1.375 s for pSOPT, and 2.4 s for HRV-ER.

### Supplementary Methods 3. Foundation-model configurations and reference representations

The primary configurations were BrainLM 13M/111M/650M, Brain-JEPA (approximately 85M parameters), Brain-Semantoks (approximately 63M), fMRI-LM VQ-contrastive/MAE/VQ-domain (approximately 684M each), and BrainHarmonix-F (approximately 89M). Parameter counts follow the configuration conventions in the manuscript and are not matched trainable-parameter budgets. Primary readouts were final-layer BrainLM classification-token (CLS) embeddings (512/768/1,280 dimensions), Brain-JEPA token mean (768), Brain-Semantoks teacher CLS (768), fMRI-LM stage-2 brain-token mean (1,024), and BrainHarmonix-F valid-token mean from the released target encoder (768). Brain-JEPA has no pretrained CLS token (12–16).

fMRI-LM inference used the released stage-2 Qwen3-0.6B-based models with brain inputs only, without text prompts. The released rank-1 language adapter was merged before benchmark adaptation. The final readout pooled the brain-token representations, rather than the stage-1 image encoder alone. Brain-Semantoks restored the teacher encoder. BrainHarmonix-F restored the encoder_ema target weights. Checkpoint identifiers and configuration specifications are provided in Table S2.

Primary downstream extraction froze encoder parameters and disabled random input masking or shuffling. The primary readout was held fixed before and after post-training. Input masks for unavailable networks or valid tokens remained active. A separate frozen-model temporal-order sensitivity deliberately shuffled observed frames as described in Supplementary Methods 14. No model-size effect was isolated: atlas, normalization, temporal coverage, architecture and tokenization also differed.

Static FC used pairwise correlations of regional BOLD signals, with upper-triangle edges Fisher transformed after clipping to ±0.999999. FC-Tucker used a rank-64 shared co-skewness basis fitted to training recordings only. For a region-centered, region-standardized $P \times T$ recording $Z$, its regional Gram matrix was $G=Z[(Z^T Z)\odot(Z^T Z)]Z^T/T^2$, where $\odot$ denotes elementwise multiplication. The leading 64 eigenvectors of the training-scan mean $G$ defined a common regional basis. Each recording was projected into that basis, and the upper-triangle Fisher-transformed connectivity among the projected signals formed the features. The basis was refitted independently in each inner and outer training partition; source-fitted coordinates were reused for transfer. Atlas-specific reference panels retained their own projection implementation; an FM score from that panel need not equal the score in the separate exact-PCA Schaefer100 comparison. Scores and paired tests must be read within their declared panel (17).

BrainGFM was evaluated as a frozen FC-based reference. The released encoder checkpoint was strictly loaded, with all encoder parameters restored and frozen. Schaefer100 Pearson matrices were supplied as node features; adjacency used FC>0.3. BrainGFM used the neutral disease token without diagnostic labels and concatenated mean/max pooling of layer-normalized final-layer node embeddings, yielding 512 dimensions. Direct Schaefer100 FC used Fisher-transformed upper-triangle edges, clipping correlations to ±0.999999 before arctanh. BrainGFM instead received the original Pearson matrices, not Fisher-transformed edges. No BrainGFM post-training was performed. Participant-level overlap with pretraining was not independently established (18).

### Supplementary Methods 4. Local prediction, clinical references, and paired-visit outcomes

Local continuous-outcome probes used ridge regression; categorical tasks used logistic regression, with binary ROC AUC or macro-averaged one-versus-rest AUC as applicable. Main nested evaluation used five outer participant-grouped folds with shuffle seeds 0, 1, and 2. The shared splitter stratified outcomes with more than ten unique values by quintiles and lower-cardinality outcomes by their observed levels. Within each outer training pool, a participant-grouped random split reserved approximately 20% for validation. The inner split was not additionally outcome-stratified except in the paired-visit arm analysis. Features were standardized to zero mean and unit variance, and principal component analysis (PCA) reduced dimensionality to limit prediction-model complexity. Both transformations were fitted separately within inner and outer training partitions and applied to the corresponding validation or test participants; validation performance selected hyperparameters without access to outer-test outcomes. In the Schaefer100 panel, regularization used C in {0.001,0.01,0.1,1,10}; ridge alpha was 1/C. Validation Spearman correlation or binary AUC selected the value. Predictions were averaged across split seeds before scoring.

Concurrent imaging-only local probes retained at most 100 PCs. Primary local HRV-ER delayed-recall probes used at most five PCs, matching the follow-up imaging budget. Primary clinical-plus-imaging follow-up models appended at most five imaging principal components (PCs) to clinical covariates. The five-component cap limited model complexity in the small intervention samples. Clinical reference variables were baseline memory, baseline EF, age, and sex in ACT, CogTE, BREATHE, and pSOPT. HRV-ER follow-up EF clinical models used baseline EF, age category, sex, and intervention arm. The primary Week-5 delayed-recall probe used imaging alone, no clinical covariates, and at most five training-fitted PCs. A separate sensitivity selected among 5, 10 and 20 PCs jointly with ridge regularization using inner validation. Historical 100-PC results are retained separately in Table S7. Main follow-up probes required the stated outcomes and clinical variables; no cognitive outcomes were imputed. Unlike paired-visit analyses, these models required baseline MRI but not follow-up MRI. Their target was subsequent cognitive level, not cognitive change or a treatment effect.

Age and sex comparisons included frozen and native-SSL-adapted encoders. MEM/EF-supervised age/sex probes were not added. HRV-ER age-category prediction was a separate exploratory multiclass task and was not interpreted as age prediction in years. Post-training diagnostic classification was a downstream probe of MEM-, EF-, or SSL-adapted representations, distinct from the earlier diagnosis-supervised training experiments. Source-exposed encoder probes were descriptive even when their local prediction heads used participant-disjoint folds.

For the HRV-ER age-category supplement, all nine frozen FM configurations, Schaefer100 FC and BrainGFM were evaluated on the same 61 baseline participants and, separately, 51 follow-up participants. The seven released categories were 54–56, 57–60, 61–63, 64–66, 67–69, 70–75 and 76–80 years. Multiclass logistic regression used the existing five-fold participant-grouped evaluation, an approximately 20% inner validation split, three split seeds and training-fitted scaling/PCA retaining at most 100 components. Regularization was selected by inner-validation balanced accuracy. Out-of-fold probability vectors were averaged across seeds before scoring. Balanced accuracy, accuracy and log loss are reported descriptively in Table S5 (Dataset S5, S5_HRV_age_category); no additional absolute-performance or paired-superiority inference was performed for this supplement.

HRV-ER follow-up EF used the same 55 participants, clinical variables (baseline EF, age category, sex and intervention arm), participant-grouped outer and inner partitions, and ridge regularization grid in two covariate encodings. For BrainLM and Brain-JEPA (†), age category and intervention arm used reference-category indicators, omitting the first category (54–56 years and the decrease arm); category columns were defined from the cohort metadata. Sex remained a binary numeric variable. For Brain-Semantoks, all three fMRI-LM configurations, BrainHarmonix-F, Schaefer100 FC and BrainGFM (‡), age category and arm used all-category one-hot indicators fitted within each training partition, with unknown categories ignored. Sex remained binary numeric. Both encodings used training-fitted standardization. The resulting clinical-only Spearman correlations were 0.730088 and 0.817965, respectively. The different coding changes the ridge penalty on category effects and can change regularization selection; the references are therefore not interchangeable. Each imaging increment was calculated against its own matched clinical reference. Table S1 provides the encoding definitions.

The same task-specific paired participants and partitions were used for all nine time-series FM configurations, Schaefer100 FC, and BrainGFM. Each visit used the same encoder checkpoint and readout. Inputs were the baseline embedding, the follow-up-minus-baseline embedding difference, or baseline concatenated with that difference. Differences were computed in float64 before fitting feature transformations. Each complete imaging input, including the concatenation, was standardized and reduced to at most five PCs in the training partition. This held total imaging dimensionality constant across the three input choices.

Regression outcomes were follow-up-minus-baseline memory or EF scores in their original cohort units; scores were not standardized separately by visit, and no responder threshold was introduced. HRV-ER had no paired memory assessment and therefore no memory-change analysis. Arm classification used four classes in ACT, three in BREATHE, and two in CogTE, pSOPT, and HRV-ER. Ridge and logistic probes used the shared C grid. Outer folds were participant-grouped and stratified by cognitive quintiles or arm labels. Inner validation reserved 20%; arm classification used an additional stratified inner split, and all arm classes were required in every partition.

Clinical change models included baseline memory, EF, age, sex, and intervention arm in ACT, CogTE, BREATHE, and pSOPT. HRV-ER used baseline EF with age category, sex, and arm. Numeric clinical covariates were imputed by training-partition medians, categorical missingness was encoded explicitly, and unseen categories were ignored at application. Clinical transformation and scaling were refitted within inner and outer training partitions. Cognitive outcomes were never imputed. Intervention-arm classification used imaging only. Null models predicted training-mean change or training class probabilities. Models incorporating follow-up MRI described longitudinal associations; they were not baseline prognosis or causal treatment-effect estimators.

Paired-visit inference retained separate families for imaging-input contrasts, adapted-versus-frozen contrasts, incremental imaging beyond clinical predictors, and frozen methods versus Schaefer100 FC. Families also separated endpoint, adaptation objective where specified, and imaging-only versus clinical-plus-imaging setting. Predictions were averaged across split seeds and then adaptation seeds before scoring. Table S8 supplies the complete primary panel. Readout extensions are in Table S4, alignment in Table S11, and input-length and PCA sensitivities in Table S7.

Table S1 (PCA_protocols) summarizes the task-specific budgets. The local delayed-recall budget was revised to five PCs for methodological consistency after historical 100-PC results were available. Frozen and adapted encoders and FC references were refitted on unchanged samples and participant partitions. Primary frozen absolute tests were repeated with 1,999 refitted label permutations; affected complete Holm families were recomputed. The dimension-selected sensitivity retains separate exploratory families and does not replace the fixed-five-PC primary result.

### Supplementary Methods 5. Cross-cohort transfer and cross-visit re-identification

The source cohort supplied the data used to fit a transferred predictor, and the target cohort was the independent cohort used for evaluation. Fixed-source cognitive axes fitted feature standardization, randomized PCA with at most 100 components, and ridge regression with alpha=10 in the source cohort and applied those transformations without target-outcome fitting. Local target-cohort probes instead fitted their prediction heads using target training participants; these two settings were reported separately. The supplied Schaefer100/BrainGFM cognitive axes retained their own source-only grouped tuning and refitting across seeds 0, 1, and 2. Their columns can be compared descriptively, but no matched significance test across different axis-fitting implementations was implied.

Intervention leave-one-study-out (LOSO) evaluation pooled the other cohorts as training data. Training outcomes were standardized separately within each training cohort; feature standardization and PCA were fitted on the pooled training participants, followed by ridge alpha=10. Held-out cohort labels were used only for evaluation. Memory LOSO used ACT, CogTE, BREATHE, and pSOPT; EF additionally used HRV-ER. A separate one-way predictor trained on ACT, CogTE, BREATHE, and pSOPT evaluated Week-5 HRV-ER recall from Week-2 MRI (57 participants). HRV-ER recall did not enter the training pools for other memory rows. These distinct endpoints were retained in the table footnotes.

Cross-visit identification used correlations between scan-level representations and nearest-match retrieval in both directions, with the gallery fixed for inference. HRV-ER used participants with both baseline and follow-up scans; NACC used its separately assembled repeated-scan sample. Paired-scan counts are reported in Table S1. Top-1 accuracy and differential identifiability (I_diff, mean matched minus unmatched similarity) were reported separately. For adapted encoders, similarities were averaged across adaptation seeds before scoring. Intervention-period pairs were not intervention-free test–retest scans. Cross-visit identification was evaluated across all seven cohorts and is shown using primary frozen readouts in the cohort radar profiles (Fig. S1) and as a difference from FC in Fig. 2; readout comparisons and additional controls are presented in Fig. S3.

Identification used fractional credit for tied best matches and averaged the two retrieval directions. Its chance reference was 1/N for a gallery of N participants. The absolute identity tests used 20,000 random bijections between the two visits, applying the same correspondence permutation in both directions. These one-sided tests assess top-1 accuracy or I_diff above the identity-permutation null; they differ from paired tests of FM versus FC. Absolute-null coverage includes the three BrainLM models, Brain-JEPA, Brain-Semantoks, the three fMRI-LM configurations, and Schaefer100 FC, with readout/projection variants as recorded. Seven primary BrainHarmonix-F top-1 tests were added to the 168 retained native-dimensional top-1 tests, giving a 175-hypothesis Holm family that includes the original alternative readouts. Other identity families, including I_diff and projected tests, were unchanged. BrainGFM has comparative identification scores but no absolute-null tests; BrainHarmonix-F has no new I_diff null tests.

The 128-dimensional identification sensitivity used fixed Gaussian projections with seeds 0, 1 and 2, shared across the two visits; similarity matrices were averaged before scoring. Native-dimensional and projected comparisons retained separate families. Their purpose was to test whether a readout contrast survived dimension matching, not to select a projection using identification outcomes.

### Supplementary Methods 6. Final-layer readouts and the focused adaptation interaction

Current readout comparisons use the same final normalized encoder output for each model. Spatial summaries average over temporal positions within the architecture's spatial units. These are 424 regions for BrainLM 13M; 27 and 31 spatial patch rows for BrainLM 111M and 650M; 450 regions for Brain-JEPA and fMRI-LM; eight observed networks for Brain-Semantoks; and 400 regions for BrainHarmonix-F. Spatial patch rows and networks are not individual anatomical ROIs. Patch overlap and valid-token weights are respected.

The spatial summaries were flattened, reduced to their feature-vector norms, or converted to embedding connectivity. For connectivity, each spatial feature vector was centered across channels and normalized to unit Euclidean norm; pairwise dot products formed a similarity matrix, whose strict upper triangle was vectorized. This differs from temporal correlation between original BOLD signals. Table S2 gives each configuration's number of units and connectivity dimension. Additional architecture-appropriate alternatives included CLS, content-token mean, CLS-plus-mean, and valid-token dispersion where supported. Downstream scaling/PCA was refitted in training partitions.

The earlier BrainLM 13M layer-3 representation is a historical intermediate-layer readout, not the current primary representation. Explicit raw layer-3 versus raw layer-4 and final-normalized-layer comparisons were kept in separate families. The final-layer alternatives were specified as a set; no single best layer/readout was selected on the held-out outcome and then reported as an independent test. Tables S4 and S7 retain the relevant multiple-comparison adjustments and matched FC controls.

An additional cognitive sensitivity selected readout within training data for all nine frozen configurations in ADNI and NACC, separately for memory and EF. It reused the original five participant-grouped outer folds and approximately 80:20 inner training/validation partitions for split seeds 0, 1, and 2. Candidates were the supported final-normalized-layer readouts: primary, flattened spatial summaries, spatial-vector norms and embedding connectivity for all configurations; content-token mean additionally for BrainLM and Brain-Semantoks; CLS-plus-mean for Brain-Semantoks; and valid-token dispersion for BrainHarmonix-F. Historical intermediate-layer and raw-layer alternatives were excluded.

Readout and ridge regularization were selected jointly by inner-validation Spearman correlation. Standardization and exact PCA (at most 100 components, subject to training rank) were fitted using inner-training participants. The ridge grid was alpha=1/C for C in {0.001, 0.01, 0.1, 1, 10}. Exact ties followed the fixed candidate order, with the primary readout first, then the fixed regularization order. After selection, transformations and the prediction head were refitted on the full outer training pool, and predictions were generated for held-out outer participants.

Selected-readout predictions reproduced the archived corresponding fixed-readout outer predictions within the specified 2 x 10^-5 numerical tolerance. Predictions were averaged over the three split seeds before scoring.

This produced 36 configuration-by-cohort-by-outcome analyses and 540 outer-fold selections. Paired references were the fixed primary readout, Schaefer100 FC, atlas-matched FC and FC-Tucker, and separately available actual-input-window FC. Table S4 (Dataset S4: S4_selected_readout_tests and S4_selected_readout_folds) contains all 164 contrasts and fold selections. This was a retrospective sensitivity analysis in previously analyzed cohorts, not a new untouched external validation. Absolute-null P values from primary readouts were not assigned to the selected pipelines.

The focused post-training-by-readout supplement reused existing checkpoints and changed only representation extraction. Its external cognitive panel covers nine configurations, two adaptation sources, MEM/EF/SSL objectives and both independent target-cohort cognitive outcomes. Primary and final-layer connectivity readouts were evaluated with target-local grouped probes; these are not fixed-source cognitive-axis transfers. Contrasts separate adapted versus frozen at the same readout, connectivity versus primary after adaptation, and comparisons with Schaefer100 FC.

This focused panel was computationally reduced after partial results were available, uniformly retaining primary and connectivity readouts rather than the original broader set. Its correction families are explicitly post hoc exploratory families. The external cognitive subset contains 540 paired comparisons in 18 families and is reported independently of the focused identification extension, which is excluded from the inferential tables. Primary-readout identification is reported separately.

### Supplementary Methods 7. Supervised and model-native self-supervised post-training

Each model was post-trained separately on ADNI and NACC for memory prediction, EF prediction, and a model-native self-supervised objective, using adaptation seeds 42, 43, and 44. The source cohort supplied post-training data; independent cohorts served as targets for external evaluation. Low-rank adaptation (LoRA) rank was 8 with alpha=16 (scale alpha/rank=2), applied to linear attention and feed-forward transformations. Matched rank did not imply equal trainable parameter counts. Supervised training minimized standardized-target mean squared error. A training-fitted StandardScaler and randomized PCA (a fixed cap of 100 components, further limited by the number of training scans minus one and feature dimensionality; random_state=0) transformed frozen features and were held fixed during adaptation. The 100-component cap was not selected by validation performance or explained variance. Downstream follow-up and paired-visit models fitted PCA separately under the five-component procedure in Supplementary Methods 4. A linear head was initialized by ridge regression with alpha=10. Training target means and standard deviations were used throughout validation and testing (19).

Encoder adapters and the prediction head were optimized jointly with AdamW at learning rate 10^-4, weight decay 0.01, effective batch size 8 (microbatch 1), and gradient-norm clipping at 1. Training lasted at most 15 epochs with patience 5. Spearman correlation on held-out source-cohort validation participants selected the checkpoint; the initial unadapted encoder and its initialized head were eligible as epoch zero. A best epoch of zero therefore denoted a valid unchanged-encoder selection, rather than an unsuccessful training result.

Within-source supervised cognition used five participant-level outer folds with an approximately 80:20 training/validation split within each training pool. Encoders, heads, target statistics, and feature transformations were fitted independently in each outer fold. Outer test participants were excluded from training and selection. Separate approximately 80:20 source participant splits selected external-evaluation checkpoints. No target-cohort outcomes entered encoder adaptation or checkpoint selection. External local probes could subsequently fit target training labels under their own grouped evaluation. Within-source summaries retained the separate aggregation recorded in Table S10 (source_cv worksheet): seed-specific metrics were supplied, while paired inference used the specified seed-averaged out-of-fold predictions.

Native SSL used source recordings with at least 200 real native frames and a shared 85:15 participant-grouped training/validation split. BrainLM reconstructed masked BOLD/image patches; reconstruction loss excluded architectural image padding for the large releases. Brain-JEPA predicted masked latent targets with its released predictor and an exponential-moving-average target encoder. Masking ratio was 75%; validation masks were held fixed across epochs. Native SSL used effective batch 8 (microbatch 1 for the original BrainLM/Brain-JEPA training loop), AdamW learning rate 10^-5, weight decay 0.05, gradient clipping at 1, at most 30 epochs, and patience 8. Loss on held-out source-cohort validation participants selected checkpoints, including epoch zero. External predictions averaged the three adaptation seeds. Source-cohort downstream probes after SSL remained source-exposed and were not independent encoder-validation results.

The added-model native SSL loops processed up to eight scans per optimizer step rather than using the original microbatch-one accumulation loop. Supervised training reused the common supervised engine. Effective batch size and source partitions were matched; the native objectives and their computation graphs were model-specific.

Brain-Semantoks SSL retained teacher–student learning with masked cosine and MCR-style representation losses, approximately 75% network-time masking, two 100-frame crops from the retained context, and an EMA teacher (0.996). A singleton training tail was duplicated where needed for the covariance regularizer. This objective is distinct from supervised cognitive regression.

fMRI-LM MAE adaptation linked the released stage-1 reconstruction decoder to the stage-2 image encoder and optimized masked BOLD reconstruction. The VQ-contrastive and VQ-domain branches used their reconstruction and quantization objectives. Decoder/quantizer parameters required by each native objective were trainable. No text-alignment objective was introduced, and the Qwen language path did not receive a separate native SSL gradient; changes in the image encoder could nevertheless change its downstream brain-token representation. Thus model-native SSL did not imply updating every component of the language-integrated architecture.

BrainHarmonix-F SSL predicted masked latent targets with the released predictor, valid-token masking and an EMA target encoder (0.996), using Smooth L1 loss. Brain-JEPA retained its released predictor/EMA framework. Architectural image padding was excluded from BrainLM reconstruction loss, which does not imply that all padded input frames were invisible to the encoder.

Table S10 supplies 162 external checkpoint-selection records (nine configurations × two sources × three objectives × three seeds) and 540 fold-specific supervised checkpoints (nine × two sources × two outcomes × three seeds × five folds). Best epoch, initial-checkpoint selection, validation score, epochs run and available trainable-parameter counts are reported. Epoch-zero selection is a valid result. Downstream analyses on the source participants used for external-checkpoint adaptation remain source-exposed, even when a later probe groups its folds by participant.

### Supplementary Methods 8. Target-to-source embedding alignment

Alignment evaluated fixed-source cognitive-axis transfer and locally fitted baseline-imaging follow-up/recall prediction without updating the encoder. Source outcome-available recordings fitted float64 StandardScaler and randomized PCA with at most 100 components; the fixed cognitive head used ridge alpha=10. The unaligned control, mean/SD matching, and correlation alignment (CORAL) shared this source fit. The unaligned control was a refitted numerical baseline for this sensitivity analysis rather than a replacement for archived primary predictions. Table S11 includes the numerical audit against those earlier predictions.

Source and calibration statistics were computed from participant-mean embeddings in source PCA coordinates. Mean/SD matching used population standard deviations floored at 10^-6. CORAL used Ledoit–Wolf covariance estimates with eigenvalues floored at max(lambda_max×10^-8,10^-10). No calibration outcomes were used. Fixed-source evaluation split target participants into five label-independent folds for seeds 0, 1, and 2; calibration used the other folds, and prediction used the held-out participants. Target calibration could include outcome-unavailable participants satisfying the specified imaging eligibility conditions (20).

Locally fitted prediction refitted calibration and subsequent target StandardScaler/PCA (at most five imaging PCs) within every relevant inner and outer training partition. Ridge alpha was selected from {0.01,0.1,1,10,100} by inner-validation Spearman correlation. Positive coordinate-wise mean/SD alignment is algebraically canceled by subsequent coordinate-wise target standardization in this pipeline. Numerical invariance was checked before reusing identical predictions; those exact identities have delta=0, CI=[0,0], and P=1. They are properties of the pipeline, not evidence that domain shift was absent. Adapter predictions were averaged across seeds 42/43/44, and Table S11 retains all scores and individual-seed summaries.

The intervention extension evaluated baseline, change and baseline-plus-change inputs for cognitive change and arm classification. Alignment was fitted using the permitted source and target training/calibration participants, then applied to held-out target participants without their outcomes. Corresponding inner training partitions were used during hyperparameter selection. Comparisons distinguished an unaligned control using the same source-PCA coordinate system from the archived primary pipeline and Schaefer100 FC. A gain over the refitted same-PCA control is not automatically a gain over the original primary model. Full controls are retained in Table S11 and Fig. S9.

### Supplementary Methods 9. Physiological outcomes and window representations

Physiological endpoints were ECG-derived heart rate in BREATHE and pSOPT and PPG-derived pulse rate in HRV-ER. They were not interchangeable with HRV/PRV variability metrics. The common window definition retained the first 160 real native fMRI frames in eight nonoverlapping 20-frame windows: 20.2 s in BREATHE, 27.5 s in pSOPT and 48 s in HRV-ER. At least six finite, time-aligned windows per scan were required. ECG rate bounds of 40–120 beats/min and BREATHE resting run-01 were retained. Usable_HR, Usable, RunUsable and related physiological usability flags were not exclusion criteria. Cognitive outcome availability was not required.

The resulting common physiological sample contained 106 participants, 187 scans and 1,494 windows in BREATHE; 36 participants, 49 scans and 386 windows in pSOPT; and 60 participants, 105 scans and 840 windows in HRV-ER. Physiological and cognitive participant counts therefore differed.

Two representation protocols were kept separate. The contextual protocol for the three BrainLM configurations and Brain-JEPA pooled final temporal tokens according to their overlap with each window while retaining the encoder's full input context. The independent-window protocol covered all nine configurations: each native window was cropped before model-specific resampling, normalization and padding, then encoded separately. Whole-recording upstream denoising still preceded this crop. Independent-window inputs can require substantial padding and are not equivalent to extracting a temporal token from a longer contextual input.

Within-scan models centered features and rates by scan and included window-position-by-visit controls; predictions were also centered over retained windows. Scan-average models averaged the same retained windows and used visit controls. Five participant-grouped outer folds and grouped 20% inner validation were repeated across seeds 0, 1 and 2, without outcome stratification. All windows and visits from a participant remained in one partition. Training-only PCA retained at most ten imaging components from the shared up-to-20-component basis; ridge regularization used the documented C grid. Same-window atlas FC and nuisance-only references were evaluated on the identical samples.

The complete nine-configuration independent-window panel reports frozen and post-trained comparisons separately in Table S9 and uses two-sided paired prediction swaps with its own Holm families. Earlier contextual, regional-amplitude and usability-filtered exports are retained as separately labeled protocols, not additional primary comparator methods. Historical HRV rate/log-RMSSD adaptation, one-sided bootstrap tests and synchronized temporal shuffling retained their original procedures. The historical 200-shift refitted circular nulls have minimum finite $P=1/201$; the retained 504-test family cannot reject at its first Holm threshold. These historical nulls are descriptive and do not establish absence of physiological information. The 1,999-shift primary frozen analysis is reported separately in Table S12 and Fig. S2.

### Supplementary Methods 10. Fusion, input context, covariates, and PCA sensitivity

Dimension-controlled fusion compared FC, FM, and their combination at equal total imaging budgets. Observational cognition used 100 PCs (100 FC, 100 FM, or 50+50), whereas clinical follow-up models used ten imaging PCs (10 FC, 10 FM, or 5+5). Clinical+FC5 was also retained to distinguish a changed FC allocation from added FM information. Transformations and tuning were fitted within training partitions. These panels used atlas-specific reference inputs and their corresponding samples, not the whole-scan Schaefer100 matrices. Complete candidate and reference scores are retained even when the corrected difference was nonsignificant.

Covariate sensitivities used only available demographic/acquisition variables, including age or age category, sex, education, site, arm, TR and real frame count. The earlier cognitive robustness panel used pSOPT baseline FD only. Separate seven-cohort FD-adjusted Re-ID and scan-average FD prediction analyses are described in Supplementary Methods 14 and 16; these did not refit the cognitive outcome analyses. ADNI one-scan sampling selected the earliest scan before outcome filtering. Session-weighted primary metrics and participant-weighted sensitivities represent different estimands. No experiment independently manipulated age, disease, TR, scanner or signal-to-noise ratio.

The 80-frame sensitivity restricted recordings to their first 80 real native frames before the model-specific resampling and input construction. Native TR was retained at cropping, so 80 frames represented different durations across cohorts. The intervention extension re-extracted both visits and repeated change/arm analyses. Shortening context also increased padding for some models; these comparisons do not isolate temporal duration from padding. The original complete-context subset required at least 200 native frames for every model. A separate nine-configuration coverage audit used the actual model-specific context requirements; both definitions remain distinguishable in Table S7.

Paired-visit PCA sensitivity jointly selected the imaging dimension from 5, 10 or 20 requested components and the prediction regularization using inner training/validation data. Rank was capped by each training partition. The final outer-training model used the selected dimension, without outer-test outcome access. Frozen and MEM/EF/SSL-adapted embeddings used the same procedure. Comparisons evaluated selected dimensionality versus fixed five components and adapted versus frozen under the selected procedure. This extension covers cognitive-change and intervention-arm tasks, not an automatic rerun of every source-axis, follow-up-level or physiological task. All 8,211 paired contrasts, scores and fold-level PC choices are in Table S7; detailed selection records are supplied in Dataset S14.

Documentary pretraining-source checks did not verify participant-level overlap.

### Supplementary Methods 11. Statistical inference, multiplicity, and consistency

Continuous tasks reported Spearman correlation and available prediction errors (MAE and RMSE; R-squared where supplied). Binary classification used ROC AUC; multiclass arms used macro-averaged one-versus-rest AUC with whole probability vectors. HRV-ER age-category classification used balanced accuracy. Other classification metrics remained supplementary. Absolute cognitive errors retained the instrument units and were not pooled across different instruments. Main paired comparisons used 5,000 participant-cluster bootstrap resamples for pointwise 95% difference intervals and 20,000 two-sided participant-level swaps of fixed predictions. Entire repeated-observation blocks were resampled or swapped together. Bootstrap ranks were recalculated with resampling multiplicities, and finite swap P values used (1+b)/(B+1), with valid draws recorded. Paired-visit multiclass swaps exchanged the full probability vector jointly.

Paired confidence intervals and prediction-swap tests conditioned on fitted out-of-fold predictions and therefore omitted variability from repeating the full model-fitting procedure. Absolute predictive tests instead refitted the prediction pipeline as described in Supplementary Methods 12. Re-ID inference additionally conditioned on the fixed set of candidate scans used for matching and used participant identification scores; adaptation ensembles averaged similarity matrices before scoring. ADNI primary metrics retained scan/session weighting while resampling participants; Table S7 provides a separate participant-weighted sensitivity. Historical HRV physiological comparisons used one-sided bootstrap tails rather than two-sided prediction swaps. Invalid draws, non-estimable contrasts, exact prediction identities, and their relevant budgets were retained in the numerical tables rather than assigned artificial zero scores.

Holm correction was applied within named exploratory families, not once across the entire benchmark. The numerical tables report correction-family membership, raw and adjusted P values, resampling counts, and uncertainty definitions for the corresponding analyses. Table S2 inventories family membership and the panel-specific aggregation rules. Family definitions differ across the primary Schaefer100, model-atlas, readout, post-training, fusion, robustness, alignment, historical, and paired-visit panels; they must not be pooled or inferred from their location within the supplementary tables. Confidence intervals were pointwise rather than multiplicity-adjusted. Nonsignificance after correction does not demonstrate equivalence or rule out smaller effects.

Primary comparison families included all nine configurations. Numerical tables identify the authoritative adjusted P value as p_adjusted and its family size as final_family_size where these fields apply. Final-layer readout, intervention readout, alignment, input-length, PCA and focused post-training-by-readout comparisons retained separately defined exploratory families. Adjusted P values were not recomputed after selecting comparisons for a figure or dividing results among tables.

New readout-selection contrasts used separate Holm families for each cognitive outcome and reference method across all nine configurations and both observational cohorts (18 tests per family for common references; 10 for the separately available actual-input-window reference). Re-ID controls used separate signal, random-initialization and motion families, each divided by metric and dimensionality mode; all model, cohort and specified contrast combinations were included within each family, or model/calibration-split/contrast combinations for motion. Temporal-order contrasts included all nine configurations, seven cohorts and two readouts within each metric-by-dimensionality family (126 tests). Table S2 (S2_readout_control_families) supplies the complete new family inventory. All new paired analyses used 5,000 participant bootstrap resamples and 20,000 two-sided swaps; these budgets are distinct from the five random initializations or five shuffled input realizations. Random and temporal-control reference scores averaged per-participant scores across five realizations, not embeddings or similarity matrices. Inference remained conditional on the fitted predictions or fixed gallery and the realized controls. The expanded FD analysis uses four seven-cohort Re-ID families of 756 contrasts and two FD-prediction families of 70 and 63 contrasts (Table S2, S2_FD_inference_families). Historical pSOPT-only motion probabilities are retained for provenance and are not the current seven-cohort correction.

For the primary frozen evidence matrix, above-chance support required a significant one-sided absolute test in at least one cohort after correction. Superiority over FC required a corrected paired advantage over the designated FC reference in at least one cohort. Cross-cohort consistency required corrected superiority to the same reference in at least two independent cohorts and positive differences in every evaluable cohort for the same task and protocol. A numerical advantage alone met none of these inferential criteria. The criteria are nonordinal: a paired FC advantage does not guarantee absolute significance. The retrospective consistency summaries for other panels retain their own task, readout, source and reference boundaries and exclude the encoder-adaptation source cohort. Table S12 provides the cohort-specific tests and supporting counts underlying Table S14; Table S2 provides the broader comparative grouping keys. The criteria were not prospectively registered.

Absolute tests and paired superiority tests address different null hypotheses and retain separate correction families. The final refitted absolute tests cover the primary frozen scope described in Methods 12, not every adapted or alternative-readout result. A nonsignificant paired difference does not establish equivalence. Pointwise bootstrap intervals and paired prediction-swap tests condition on fitted predictions; the label-permutation tests instead refit the specified prediction pipeline.

Figure 2 summarizes primary frozen performance in a cohort-grouped heatmap. Color is the archived method score minus matched Schaefer100 FC, in the task-specific units of Spearman correlation, AUC, balanced accuracy for HRV-ER age-category classification, or Re-ID accuracy. Warm colors indicate higher performance than Schaefer100 FC, cool colors indicate lower performance, and white indicates zero difference. Hatching denotes unavailable outcomes. Statistical significance is not marked in this figure. The FC reference row has zero color difference by construction; its vertically arranged numbers report the actual FC scores, rounded to two decimal places. A positive color alone does not establish a significant paired advantage. Score differences are not pooled across metrics. Follow-up cognition uses baseline imaging alone. ΔCOG and ARM used imaging-only follow-up-minus-baseline inputs: differences between separately extracted primary frozen embeddings for the FMs and BrainGFM, and between FC edge vectors for Schaefer100 FC. Differences were computed before training-fitted standardization and PCA (at most five components). HRV-ER memory follow-up retains its single delayed-recall endpoint and five-PC protocol. The heatmap additionally includes intervention demographic and baseline-cognition tasks from main-text Table 2. HRV-ER age is evaluated as seven-category classification (balanced accuracy); baseline memory and memory-change outcomes are unavailable. Scores and test provenance are provided with Fig. 2 and in Dataset S15. As in Supplementary Methods 12, archived paired-prediction scores are retained while absolute tests use their specified refitting procedures.

### Supplementary Methods 12. Absolute tests and evidence matrix

Absolute predictive tests covered the nine primary frozen configurations. Within-cohort age, sex, diagnosis, memory and EF tests were limited to ADNI and NACC. Intervention tasks comprised imaging-only follow-up memory and EF, associations with memory and EF change using follow-up-minus-baseline representations, and follow-up-minus-baseline intervention-arm classification. HRV-ER single delayed recall joined the follow-up-memory display and correction family, while retaining its distinct outcome and eligibility, with the same at-most-five-PC budget as the other primary follow-up probes. Other imaging-only follow-up probes retained the corresponding clinical-panel eligibility but removed clinical predictors and used at most five imaging PCs. Fixed-source transfer, LOSO, adapted encoders, alternative readouts, baseline/combined change inputs and scan-average physiology were outside the absolute-test scope.

For each predictive hypothesis, 1,999 label permutations were evaluated with one-sided P=(1+b)/(B+1), where b counts null scores at least as large as the observed score. Three split seeds (0, 1, 2) and participant-grouped five-fold outer evaluation were retained. Outcome-class or continuous-outcome-quintile stratification was regenerated for each permuted outcome. Grouped approximately 20% inner validation was regenerated, with arm stratification for paired-visit classification. Scaling, PCA, regularization selection and ridge/logistic prediction were refitted within training partitions; held-out predictions were averaged across seeds before scoring. ADNI outcome trajectories were permuted as whole participant blocks only between participants with the same retained visit count. All visits stayed grouped; session-level scoring weights were retained. This exchangeability restriction does not remove all potential cohort or acquisition confounding.

Holm correction used one family per predictive task across all nine configurations and eligible cohorts: 18 tests each for age, sex, diagnosis, memory and EF; 45 each for follow-up memory, follow-up EF, EF change and arm classification; and 36 for memory change, which excluded HRV-ER. A uniform budget of 1,999 permutations was used for computational feasibility, without significance-dependent stopping. The minimum raw P for 1,999 permutations is 0.0005.

Independent-window within-scan physiological tests used 1,999 independent scan-wise circular shifts of the rate series, including possible zero shifts, and a 27-test Holm family (nine configurations by three cohorts). Shifts retained within-scan serial ordering up to the circular origin and assumed cyclic exchangeability. Prediction models and outcome-dependent tuning were refitted. Outcome-independent grouped partitions and unsupervised training-partition transformations could be cached without changing the null procedure. The pipeline included window-position-by-visit nuisance terms, scan centering and the frozen window embeddings. Significance therefore supported temporal alignment of the complete procedure, not incremental embedding value, neural specificity or a causal dependence on temporal ordering. Paired nuisance/FC comparisons remained separate.

Identity tests used 20,000 random participant-identity bijections and one-sided finite permutation probabilities, preserving the same mapping in both retrieval directions and fractional ties. The enlarged native-dimensional top-1 family contained 175 hypotheses; the 168 retained tests were not reduced to primary readouts before adding seven BrainHarmonix-F tests. The matrix used primary top-1 only. Differential identifiability and other historical identity families retained their original inference.

Observed and null predictive scores were obtained from the same CPU refitting implementation. Continuous-outcome reproduction was checked against archived predictions with a 2×10^-5 numerical tolerance. In classification, validation-AUC ties and solver roundoff could change the selected regularization and predictions; aggregate discrepancies are supplied in Table S12. Primary prediction scores and paired inferences were retained; refitted observations are reported with their corresponding absolute-null P values. Matched imaging-only follow-up FC/BrainGFM comparisons used 5,000 participant bootstraps and 20,000 two-sided prediction swaps, with four task-by-reference Holm families of 45 tests.

Table S14 displays corrected absolute-test evidence separately for each cohort: ● indicates significant above-chance performance, ○ indicates no significant above-chance evidence, and – indicates an unavailable outcome. Symbol positions follow the cohort order in the table footnote. Superiority over FC is assessed separately using corrected paired comparisons and is not encoded by these symbols. Dataset S12 additionally supplies the cohort-specific evidence display (S12_cohort_evidence_matrix) and its cohort-specific adjusted tests (S12_cohort_evidence_tests). The supplementary cohort-aggregated worksheet S12_evidence_matrix uses a different display: ✓ denotes superiority over FC in at least one cohort, ○ denotes absolute support without demonstrated superiority over FC, and × denotes neither criterion being supported after correction. This supplementary summary uses different symbols from Table S14. Corrections were applied over the complete families before aggregation. The criteria are nonordinal: superiority over FC does not imply significant absolute performance. Neither an open circle in the supplementary evidence table nor a cross in the supplementary summary establishes absence of signal or equivalence to chance. Re-ID uses top-1 accuracy; supplementary physiology evidence uses the within-scan independent-window protocol. Schaefer100 FC is the reference for scan-level tasks, and corresponding same-window atlas FC is the physiological reference. No configuration met the specified cross-cohort FC-superiority consistency criterion. Full cohort-specific tests, denominators, paired reference results, and supporting counts are supplied in Table S12. No participant predictions or individual clinical records are included in the supplementary datasets.

### Supplementary Methods 13. Data, code, and supporting materials

Individual-level data are governed by the cohort-specific access conditions described in the main manuscript. Access to ACT, CogTE, BREATHE and pSOPT data may be requested from the Stanford CogT lab through Feng Vankee-Lin, subject to applicable institutional approvals and data-use agreements. Custom code is maintained at https://github.com/ymliu17/brainfm.

This appendix and the accompanying datasets provide aggregate results, supplementary figure source data, model specifications and analysis definitions. Individual outcomes, participant-level predictions and pretrained weights are not redistributed. Original pretrained checkpoints were obtained from the model developers. Requests for post-trained adapters should be directed to the corresponding author and are subject to applicable data-use conditions.

### Supplementary Methods 14. Additional Re-ID controls

Re-ID controls covered all nine frozen configurations and seven cohorts, using identical matched scan pairs and model-specific input construction. Paired galleries comprised 435 participants in ADNI, 116 in NACC, 95 in ACT, 73 in CogTE, 107 in BREATHE, 43 in pSOPT and 51 in HRV-ER. Primary and embedding-connectivity readouts used the same final normalized model output. Native-dimensional and fixed 128-dimensional Gaussian-projection comparisons followed the same tie-aware bidirectional scoring and differential-identifiability procedures as the existing readout panel.

Five random initializations (seeds 101–105) were evaluated without training, with each initialization shared across visits and cohorts. Learned encoder components were initialized using the official architecture constructors while fixed atlas, gradient and positional priors were retained. For fMRI-LM, both the learned tokenizer and language backbone were randomized; the contrast does not separate imaging and language pretraining contributions. Existing BrainLM 13M random features were reused after verifying scan pairs and input provenance. Pretrained extraction was checked against the archived features before random-model extraction. The original canonical batches of four were retained for fMRI-LM to avoid batch-dependent bfloat16 differences; other configurations used single-scan batches.

Signal references were regional BOLD standard deviations (population normalization, ddof=0), lag-one autocorrelations, and input-window-matched FC. Summaries used real frames within the model-matched retained windows, excluding temporal padding and unavailable regions. For the original BrainLM/Brain-JEPA configurations, canonical time-series inputs were used. For added configurations, verified source time series followed the model-specific region ordering and resampling before summary extraction, but preceded model-specific regional scaling. Standard-deviation controls can therefore retain amplitude information removed by model normalization. They were evaluated at native dimensionality and under the common 128-dimensional projection.

Random-model reference scores averaged participant-level scores across the five initializations. Contrasts compared pretrained and random models at each readout and tested the pretraining-by-readout interaction: the pretrained connectivity-minus-primary gain minus the mean corresponding random-model gain. Signal contrasts compared primary and connectivity readouts with standard deviation and lag-one summaries, and connectivity with input-window-matched FC. The original control release contained 2,448 contrasts: 1,260 signal, 756 random and 432 pSOPT-only motion comparisons. The expanded seven-cohort motion analysis contains 3,024 contrasts and supersedes the 432 motion rows for current FD inference; signal and random families are unchanged. Holm families contained 315 signal or 189 random contrasts per metric and dimensionality mode. These nine-configuration families supersede the earlier 13M-only control inference for the expanded panel; historical scores and original P values remain explicitly labeled in Dataset S13.

Paired scan-average framewise displacement (FD) was available for 435 participants in ADNI, 116 in NACC, 95 in ACT, 70 in CogTE, 107 in BREATHE, 43 in pSOPT and 51 in HRV-ER. Three CogTE participants lacked paired FD and were excluded from this sensitivity only. FD was joined by participant and visit: ACT baseline/follow-up corresponded to sessions 01/03; CogTE used T1/T2; BREATHE used resting-state run-01; HRV-ER used pre/post mean FD. No additional FD-threshold exclusions or FD imputation were applied to the retained benchmark sample beyond the upstream quality control described in Supplementary Methods 2. Coverage is given in Table S1.

For each configuration, ordinary least squares with an intercept and log(1+FD) was fitted jointly to both visits of 20% calibration participants, separately for each representation, and applied to the remaining 80%. Three participant splits used seeds 0, 1 and 2. Evaluation participants and galleries were identical across adjusted methods and the corresponding unadjusted control. No encoder weights were updated. Native-dimensional and fixed 128-dimensional projection analyses used the existing Re-ID scoring protocol. The overlapping calibration splits are sensitivity analyses, not independent replications.

Contrasts compared adjusted embedding connectivity with adjusted primary representations, adjusted input-window FC, adjusted random-connectivity scores averaged over five initializations, and unadjusted pretrained connectivity on the same gallery. The seven-cohort analysis included 3,024 contrasts, with four separate Holm families defined by metric and dimensionality; each family contained 9 configurations × 7 cohorts × 3 splits × 4 contrasts = 756 tests. Paired inference used 5,000 participant bootstraps and 20,000 two-sided participant-score swaps conditional on the fitted adjustment, fixed gallery and realized random controls. Previously computed pSOPT scores and raw P values were reused after checking FD and split agreement; their adjusted P values were recalculated in the expanded families.

Mean FD described the whole scan and was not necessarily matched to the truncated model input. Residualization included an intercept and one linear log-FD term; persistence of a readout advantage does not establish independence from all motion effects. Comparisons of adjusted and unadjusted connectivity include the consequences of removing the fitted intercept as well as the FD association.

Temporal-order sensitivity used five shuffled realizations per scan (seeds 211–215). Within each scan, a common permutation was applied across observed regions to the real frames consumed by the encoder, after model-specific input construction and before tokenization. Visits used independent scan-specific permutations. Padding, unavailable-input rows, validity masks and acquisition metadata were unchanged. This operation preserved static FC and regional signal distributions. Implementation checks verified a common bijective permutation and unchanged excluded input positions; numerical checks of FC, means and standard deviations confirmed invariance.

Frozen primary and final-layer embedding-connectivity readouts were evaluated for original and shuffled inputs using the same seven galleries, at native dimensionality and after projection to 128 dimensions. The shuffled reference averaged participant-level scores across the five realizations. The 504 paired contrasts tested original minus shuffled accuracy or differential identifiability, with the 5,000-bootstrap and 20,000-score-swap procedure and 126-test Holm families described in Supplementary Methods 11. Five shuffled realizations were perturbation repeats, not an absolute-null permutation test. A difference indicates sensitivity to this input perturbation; it does not establish neural specificity, a causal temporal mechanism or superiority over FC. This panel did not test cognitive or physiological outcomes.

Table S13 (Dataset S13) supplies the complete control, temporal-order and motion analyses. S13_nine_config_controls remains authoritative for signal and random panels; its pSOPT-only motion rows are historical. S13_FD_ReID_tests supplies the current seven-cohort FD correction, and S13_temporal_order retains the temporal-order results. Historical13_* worksheets retain the earlier BrainLM 13M-only inference. S13_FD_prediction_scores and S13_FD_prediction_tests supply the separate scan-average FD regression analyses described in Supplementary Methods 16.

### Supplementary Methods 15. Regional signal variability beyond Re-ID

Regional BOLD SD was evaluated as a supplementary signal control for frozen primary readouts and final-layer embedding connectivity across all nine configurations. These analyses supported interpretation of the readout findings and were separate from the primary benchmark comparisons with FC-based representations and clinical predictors. The extension covered ADNI and NACC memory and EF, baseline cognitive scores in ACT, CogTE, BREATHE, pSOPT and HRV-ER, and intervention follow-up cognition, cognitive-change associations and intervention-arm classification. HRV-ER had baseline and follow-up EF and a single delayed-recall outcome predicted from baseline imaging; baseline memory and memory change were unavailable. The extension did not rerun post-training, cross-cohort transfer or alignment combinations. It was an exploratory analysis motivated by the completed Re-ID controls.

SD was calculated across retained real frames for each region, using population normalization (ddof=0), excluding temporal padding and unavailable cerebellar regions. Five input families were shared across the nine configurations: A424, first 200 native frames (three BrainLM models); Schaefer400+Tian50, first 160 native frames (Brain-JEPA); Schaefer400+Tian50, first 100 or 160 frames after resampling to TR=2 s (Brain-Semantoks or fMRI-LM, respectively); and Schaefer400, up to 18 patches of round(35.28/TR) native frames (BrainHarmonix-F). Short recordings contributed only observed samples. SD preceded model-specific regional scaling and could retain amplitude information removed by those transformations. SD values were verified to agree exactly with the completed Re-ID controls on overlapping scans; the prediction panel also included eligible participants without paired Re-ID scans.

Local cognitive-level probes retained the existing task-specific scan IDs, labels, participant-grouped outer folds and inner validation partitions. Scaling and PCA were fitted in each training partition, retaining at most 100 components for cognitive-level prediction and five for follow-up and delayed recall. Ridge regularization used the existing grid and validation procedure, with predictions averaged across split seeds 0, 1 and 2. Follow-up analyses used baseline SD, with imaging-only and clinical-plus-imaging settings. The separate HRV-ER clinical delayed-recall panel retained its existing clinical-block probe, using baseline EF, age category, sex and arm, with at most five imaging components. No baseline memory value was imputed.

Paired-visit analyses used baseline SD, follow-up-minus-baseline SD, or concatenated baseline and change SD, with one training-fitted PCA retaining at most five imaging components. Cognitive-change analyses included imaging-only and clinical-plus-imaging models; intervention-arm classification used imaging only. The same paired-visit samples, class order, outer and inner splits, covariate transformations and ridge/logistic tuning were retained. These analyses involving follow-up MRI assess longitudinal associations and arm classification, not pretreatment prediction of benefit.

FM references used frozen primary and final-layer embedding-connectivity readouts. Completed predictions were reused when samples and protocols matched; otherwise the existing prediction procedure was applied to cached representations. Local FC references included Schaefer100 FC and FC from the same model-input window. Paired-visit FC comparisons used the existing Schaefer100 FC results. No encoder was updated or re-extracted. Original prediction results and correction families were preserved.

The panel contained 2,696 paired comparisons. Each used 5,000 participant bootstrap resamples and 20,000 two-sided swaps of fixed predictions, retaining participant blocks for repeated scans and swapping entire probability vectors for multiclass arm outcomes. Confidence intervals were pointwise and conditional on fitted predictions. Local Holm families were defined by endpoint, imaging-only versus clinical-plus-imaging setting, and contrast class (FM versus SD, FC versus SD, or clinical-plus-SD versus clinical). Each family pooled applicable configurations/readouts, input families and cohorts. Longitudinal families additionally pooled all three input modes within the same endpoint, setting and contrast class. HRV-ER delayed recall contributed to the corresponding follow-up-memory family while retaining its distinct outcome definition. No SD absolute-performance permutation tests were added; relative superiority must not be interpreted as above-chance evidence. Table S15 (Dataset S16) provides all family sizes and contrasts.

### Supplementary Methods 16. Prediction of scan-average framewise displacement

Concurrent scan-average FD was predicted separately in seven cohorts from the nine frozen primary scan-level representations, frozen BrainGFM, Schaefer100 FC, and five model-input-matched regional SD controls. ADNI retained available repeated scans; NACC used the selected single scan per participant; intervention cohorts included available pre/post scans. BREATHE included its nonrandomized subgroup for this technical outcome, and HRV-ER retained the older-adult subgroup. Samples were intersected across all 16 methods within each cohort. Analysis counts (scans/participants) were ADNI 1,651/712, NACC 560/560, ACT 190/95, CogTE 141/71, BREATHE 214/107, pSOPT 87/44 and HRV-ER 112/61. No additional FD threshold was applied to the retained benchmark sample beyond upstream quality control. Table S1 distinguishes missing FD from missing comparator features.

Prediction used ridge regression with log(1+FD) as the target. Five outer folds grouped all scans from a participant and were stratified by FD quintiles. Each outer training pool used an approximately 80:20 participant-grouped random training/validation split. Seeds 0, 1 and 2 defined three repeated evaluations, shared by all methods. Standardization and exact PCA, retaining at most 100 components subject to training rank, were fitted within inner-training partitions. Ridge regularization was selected by validation Spearman correlation from alpha=1/C, C in {0.001, 0.01, 0.1, 1, 10}. Transformations and regression were refitted on each outer training pool before predicting its held-out participants.

Out-of-fold log-FD predictions were averaged across split seeds and transformed back with expm1. Spearman correlation used untruncated predictions; negative predicted FD values were clipped to zero only for mean absolute error and root-mean-square error in millimeters. A training-mean FD reference supplied error benchmarks. Scores retained scan-level weighting while uncertainty resampled participants. Pointwise 95% correlation intervals used 5,000 participant-cluster bootstraps conditional on fitted predictions.

Paired inference used 5,000 participant bootstraps and 20,000 two-sided participant prediction swaps without refitting. Holm correction used separate families of 70 comparisons against Schaefer100 FC (nine configurations plus BrainGFM across seven cohorts) and 63 comparisons against the model-input-matched SD control (nine configurations across seven cohorts). No refitted absolute-null tests were performed; correlation intervals were not interpreted as corrected above-chance evidence. The analysis assessed motion-related information in representations, not the fraction of cognitive or identification performance attributable to motion. Full scores, errors, intervals and comparisons are in Table S13.

### Supplementary Results

#### 1. Concurrent associations and reference comparisons

Tables S3 and S5 provide complete numerical scores and paired comparisons, including nonsignificant effects. Primary Schaefer100 comparisons and atlas-/input-matched comparisons are separate panels. In ADNI, Schaefer100 FC concurrent-memory correlation was 0.403, compared with a maximum of 0.284 among the nine primary FM configurations. In NACC, fMRI-LM MAE concurrent-memory correlation was 0.295 and Schaefer100 FC was 0.194; the numerical advantage did not survive the declared correction. NACC all-cause dementia discrimination ranged from AUC 0.793 to 0.858 across FMs, versus 0.828 for Schaefer100 FC. The smaller AD-etiology analysis is retained with its own sample and diagnostic definition.

HRV-ER delayed recall is distinguished from baseline memory composites. Local baseline-imaging prediction, fixed-source cognitive axes and one-way intervention-cohort transfer remain separate analyses. Under the five-PC primary protocol, no frozen configuration outperformed its FC references after correction. The previous fMRI-LM MAE advantage over Schaefer450 FC-Tucker arose under the historical 100-PC protocol and is retained only in the archived sensitivity results. Table S3 contains the revised primary comparisons; Table S7 retains the historical contrasts.

#### 2. Cross-cohort transfer

Table S6 includes observational-source cognitive axes, intervention LOSO and the separate 57-participant HRV-ER delayed-recall transfer. Source-trained axes do not use target outcomes to fit a head; target-local post-training probes do. The LOSO memory training pool excludes HRV-ER because its single delayed-recall measurement is not a repeated MEM endpoint. Source and target dependence is reported without attributing it to a particular population or acquisition difference.

#### 3. Longitudinal inputs and PCA sensitivity

Table S8 and Fig. S7 report baseline, follow-up-minus-baseline and combined inputs across the available intervention outcomes. The nine-configuration primary families retained an HRV-ER BrainLM 111M intervention-arm advantage for change versus baseline imaging (AUC 0.729 versus 0.360), but no corresponding corrected cognitive-change input advantage. Follow-up cognitive prediction and cognitive-change associations used separate outcomes and matched clinical reference models.

Expanded PCA sensitivity retained all frozen/post-trained paired-visit scores and 8,211 paired contrasts in Table S7. The requested dimensionality was selected within inner training/validation data and capped by training rank. Its complete positive, negative and nonsignificant contrasts are supplied; a result obtained after inner dimension selection is not substituted for a fixed-five-PC primary result. The PC-selection records permit assessment of whether the five-component constraint was active and which larger budgets were chosen.

The local HRV-ER delayed-recall extension compared fixed-five-PC and inner-selected 5/10/20-PC probes for all nine frozen configurations and all six post-training source/objective combinations, separately by scan timing. Among the 126 dimension-selection contrasts, 0 showed a corrected improvement. Full scores, PC choices and archived 100-PC results are in Table S7. Baseline-MRI prediction and post-assessment MRI associations remain separate analyses.

**4. Re-ID and physiology**

Primary Re-ID accuracy is displayed in Fig. S1 and Fig. S4A; differential identifiability is displayed separately in Fig. S4B. Brain-Semantoks and BrainLM 111M had corrected positive differential identifiability across all seven cohorts in the available absolute-null panel. Identification accuracy and differential identifiability are distinct measures. The absence of a primary FM advantage over Schaefer100 FC does not negate above-chance identity information. Available absolute-null tests, paired comparisons and denominators appear separately in Table S5.

Independent-window physiological results for all nine configurations appear in Table S9 and Fig. S11. In HRV-ER scan-average pulse-rate prediction, Brain-Semantoks and fMRI-LM VQ-domain correlations of 0.164 and 0.177 exceeded the nuisance reference of −0.349 after correction, but not their FC references. Positive within-scan correlations were not interpreted as corrected improvements when the paired tests against nuisance or FC were nonsignificant. Contextual four-configuration and historical variability analyses retain their own sample and inference labels.

**5. Readout sensitivity**

Table S4 includes final-layer readouts for cognition, demographics, diagnosis, identification, cross-cohort axes and paired-visit intervention outcomes, with matched FC controls. BrainLM 13M final spatial flattening improved selected ADNI age prediction, sex and diagnostic classification, and concurrent cognitive associations relative to final CLS. Embedding-connectivity identification contrasts were also assessed after projection to 128 dimensions. These comparisons isolate readout changes more closely than the older layer-3 versus final-CLS contrast, while spatial-unit definitions still differ across architectures. Additional nine-configuration controls are supplied in Table S13.

Training-partition readout selection significantly improved five of the 36 configuration/cohort/outcome analyses relative to the fixed primary readout (Table S4). All five occurred in ADNI: BrainLM 13M memory (rho=0.185 to 0.390) and EF (0.133 to 0.322), Brain-Semantoks memory (0.202 to 0.258), and fMRI-LM MAE memory (-0.039 to 0.153) and EF (-0.019 to 0.110); each had Holm-adjusted P=0.0009. For concurrent memory associations in NACC, the selected fMRI-LM MAE pipeline reached rho=0.338 versus 0.194 for Schaefer100 FC (adjusted P=0.022), but did not significantly outperform its primary readout, atlas-matched FC, FC-Tucker or actual-input-window FC. No new absolute-null significance was inferred for selected pipelines.

Under the expanded nine-configuration control families, pretrained embedding connectivity exceeded the mean random-initialization accuracy in six, seven and five cohorts for BrainLM 13M, 111M and 650M, respectively, at native dimensionality. After projection to 128 dimensions, the corresponding counts were six, five and five. These results refer to paired pretraining contrasts, not absolute-null tests. No configuration significantly exceeded regional BOLD standard deviations for Re-ID in any cohort; this reference exceeded connectivity in 52 of 63 configuration/cohort comparisons at each dimensionality setting. The expanded seven-cohort motion analysis is reported in Supplementary Results 10 and Table S13.

Synchronous time shuffling reduced Re-ID accuracy in selected configurations and cohorts (Table S13). For BrainLM 13M connectivity, original versus mean shuffled accuracy was 66.3% versus 39.1% in pSOPT (adjusted P=0.017) and 38.2% versus 10.6% in HRV-ER (adjusted P=0.006); both contrasts remained significant at 128 dimensions. At native dimensionality, original-order advantages also occurred for Brain-Semantoks primary readouts in BREATHE, pSOPT and HRV-ER; fMRI-LM VQ-contrastive primary readout in NACC; VQ-contrastive and MAE connectivity in NACC, pSOPT and HRV-ER; and VQ-domain connectivity in HRV-ER. Conversely, Brain-JEPA connectivity in ACT improved after shuffling from zero to 2.0% accuracy. All contrasts, including nonsignificant and reversed effects, are retained. These findings establish sensitivity to the specified perturbation in some settings without isolating a neural temporal mechanism.

Post-training-by-readout results are reported separately in Table S10 and Fig. S8 because they reuse adapted checkpoints. Frozen readout gains do not imply that encoder adaptation is beneficial. Full intermediate-layer historical comparisons are retained with their original inference rather than silently renamed final-layer results.

#### 6. Post-training and checkpoint selection

Table S10 distinguishes independent fold-specific source evaluation, external target-local prediction, fixed-source transfer and source-exposed descriptive tests. The external-checkpoint and CV checkpoint audit includes epoch-zero selections, which indicate that adaptation did not improve the source selection metric. The primary post-training comparisons retain all three adaptation seeds and the full declared families. Main-text Fig. 4A–D shows the two corrected within-source memory gains and the two independent-cohort gains. The focused target-local readout analysis below addresses a different set of contrasts.

The focused external cognitive supplement used prediction models fitted locally in the independent target cohort and contains 216 same-readout adapted-versus-frozen comparisons, 108 connectivity-versus-primary comparisons and 216 comparisons with Schaefer100 FC. These are distinct from the fixed-source transfer analyses. None of the 216 same-readout adaptation contrasts was significant after its retained correction. Some connectivity-versus-primary comparisons were significant, but these test extraction choice after adaptation, not an adaptation benefit at fixed readout. The reduced families were defined post hoc for this computationally focused supplement. The focused identification extension is outside the reported inferential scope.

#### 7. Alignment and input context

Table S11 and Fig. S9 separate gains over the same-coordinate unaligned control from gains over the original primary pipeline or FC. Selected cognitive and intervention-arm gains were not consistent across all settings. Mean/SD matching can cancel exactly when followed by target coordinate standardization; those identities are reported analytically rather than interpreted as evidence against domain shift.

Table S7 and Fig. S10 report both favorable and unfavorable 80-native-frame effects. The contrast changes duration and, for some models, padding simultaneously. Complete-context, demographic/acquisition adjustment and participant-weighting analyses do not identify independent causal effects of acquisition parameters.

#### 8. Absolute-test evidence and comparison with reference superiority

The primary absolute-test panel comprised 333 predictive or circular-shift tests with 1,999 permutations each. Seven BrainHarmonix-F identity tests, using 20,000 mappings each, were included with the retained identity tests. Table S12 reports complete families and Fig. S2 displays corrected support. In NACC, all nine configurations showed supported diagnostic discrimination and concurrent memory associations (adjusted P=0.009), whereas none showed supported concurrent EF associations. In ADNI, five configurations each showed supported diagnosis, memory and EF prediction. Supported age prediction occurred in four configurations in ADNI and one in NACC; supported sex classification occurred in three and eight, respectively. These absolute tests do not overturn the separate paired comparisons against FC.

None of the primary imaging-only follow-up, follow-up-minus-baseline cognitive-change or follow-up-minus-baseline arm tests survived its absolute-test correction. HRV-ER BrainLM 111M change-based arm classification (AUC=0.729) had raw P=0.002 and adjusted P=0.090. HRV-ER local delayed-recall tests used the five-PC protocol; all nine were nonsignificant after correction. Absolute null tests assess prediction against shuffled outcomes; paired comparisons assess differences between methods or input constructions. Historical 100-PC delayed-recall statistics are retained separately in Tables S7 and S12.

In matched imaging-only follow-up comparisons, Brain-Semantoks predicted ACT follow-up memory with rho=0.294 versus -0.123 for Schaefer100 FC (difference=0.417; paired Holm-adjusted P=0.018). Its absolute label-permutation result was not significant (adjusted P=0.110). Thus relative superiority and absolute support did not imply one another. This result concerns imaging-only follow-up prediction, not improvement beyond baseline clinical measures.

Within the 175-hypothesis native top-1 family, Brain-Semantoks exceeded chance in six cohorts, BrainLM 111M in four and BrainLM 650M in three. BrainLM 650M did not exceed chance after correction in pSOPT (adjusted

P=0.0534). BrainHarmonix-F showed supported top-1 identification in ADNI only. Historical differential-identifiability and paired readout tests retained their original inference.

All nine configurations in BREATHE and HRV-ER showed significant within-scan physiological temporal alignment (18 of 27 tests; adjusted P=0.0135); no pSOPT test survived correction. These tests concern the full covariate-plus-embedding procedure and do not establish a corrected incremental benefit over nuisance-only or FC reference prediction. The primary frozen evidence matrix showed no configuration satisfying the specified cross-cohort FC-superiority consistency criterion. That matrix does not summarize alternative-readout or post-training benefits.

#### 9. Regional signal variability beyond Re-ID

The 1,836 FM-versus-SD contrasts yielded eight corrected advantages for FM representations and 18 for SD. These counts refer to correlated configuration, readout, input and cohort comparisons, not independent replications. For concurrent memory associations in NACC, fMRI-LM MAE embedding connectivity exceeded its input-matched SD reference (rho=0.352 versus 0.188; difference=0.164, pointwise conditional 95% CI, 0.085–0.246; Holm-adjusted P=0.020). In ADNI, SD exceeded both primary and connectivity Brain-JEPA representations for memory and EF; additional ADNI contrasts favored SD over several fMRI-LM representations. Complete results, including nonsignificant differences, are in Table S15.

Selected imaging-only follow-up comparisons favored BrainLM 650M in CogTE EF, BrainHarmonix-F in CogTE memory, and fMRI-LM VQ-contrastive/VQ-domain connectivity in HRV-ER delayed recall. Conversely, SD exceeded primary BrainHarmonix-F for ACT follow-up memory and its connectivity readout for HRV-ER follow-up EF. Several FM advantages involved negatively correlated SD references and do not establish absolute predictive validity. In the paired-visit panel, Brain-JEPA primary representations exceeded SD for ACT baseline-input arm classification (AUC=0.520 versus 0.276; adjusted P=0.0135), and Brain-JEPA connectivity changes exceeded SD changes in analyses of HRV-ER EF-change associations (rho=0.217 versus -0.478; adjusted P=0.0135). No other FM-versus-SD cognitive-change or arm contrast was significant after correction.

None of the 185 clinical-plus-SD versus clinical-only comparisons significantly improved follow-up cognitive prediction or cognitive-change associations after correction. Likewise, no FM-versus-SD contrast in the clinical-plus-imaging settings was significant after correction. SD therefore provided a useful reference for assessing task-dependent relative performance, without establishing added value beyond the available clinical predictors.

#### 10. Motion adjustment and prediction of scan-average FD

After FD adjustment, BrainLM 13M embedding connectivity exceeded its primary CLS readout in ADNI, NACC, ACT, CogTE, BREATHE and pSOPT across all three calibration splits at both native and matched dimensionality; HRV-ER differences were positive but did not survive correction. In ADNI, native-dimensional adjusted connectivity accuracy ranged from 42.2% to 46.0%, compared with 2.4%–2.6% for adjusted CLS and 31.5%–34.9% for unadjusted connectivity on the same evaluation galleries. Its adjusted-minus-unadjusted improvements were significant in all three splits (adjusted P=0.0378). At 128 dimensions, adjusted connectivity exceeded the adjusted random-connectivity reference across all three splits in the same six cohorts. Native-dimensional pSOPT random-reference superiority survived in two of three splits under the expanded correction. Fig. S3F illustrates the pSOPT subset using the updated correction; the splits are not independent replications.

Readout and pretraining effects varied across the remaining configurations. Across the 189 model/cohort/split comparisons at each dimensionality, adjusted connectivity exceeded the adjusted primary readout in 38 native-dimensional and 36 projected comparisons; one native-dimensional difference favored the primary readout. Pretrained connectivity exceeded the adjusted random reference in 39 native and 37 projected comparisons, while 8 native and 4 projected Brain-JEPA comparisons favored random initialization. No FD-adjusted connectivity accuracy significantly exceeded the corresponding adjusted input-window FC reference. All contrasts and nonsignificant results are retained in Table S13.

Correlations between predicted and observed scan-average FD varied across configurations and cohorts. In ADNI, BrainLM 13M, 111M and 650M correlations were 0.370, 0.498 and 0.534, respectively, compared with 0.673 for Schaefer100 FC. In HRV-ER, the three fMRI-LM correlations ranged from 0.621 to 0.701, compared with 0.591 for FC. All nine configurations had lower FD-prediction correlations than FC in ADNI after paired correction, and none significantly exceeded FC in any cohort. Brain-Semantoks and the three fMRI-LM configurations exceeded their matched SD controls in NACC and HRV-ER after correction. These comparisons concern motion prediction, not superiority on a cognitive outcome; absolute above-chance significance was not tested. Together, the FD analyses document motion-related information while showing that the principal BrainLM 13M Re-ID readout advantage persisted under the specified adjustment.

### Supplementary tables (external Excel files)

Tables S1–S13 are supplied as Datasets S1–S13, respectively. Detailed fold-level PCA selections accompanying Table S7 are in Dataset S14. Source data for main Figs. 2–4 and Figs. S1–S11 are in Dataset S15. Each workbook includes worksheet descriptions and field definitions. Current inferential worksheets identify the authoritative adjusted probability as p_adjusted; historical analyses retain their explicitly labeled original inference. Dataset S12 distinguishes the cohort-specific Table S14 display from the retained supplementary cohort-aggregated summaries. Table S14 reproduces the cohort-specific evidence matrix in this appendix; its numerical support is in Dataset S12. Dataset S14 remains the separate PCA-selection dataset. Table S15 is supplied separately as Dataset S16; its paired Holm-adjusted probabilities are p_adjusted (also retained as holm_p).

**Table S1. Cohorts, outcomes, acquisition metadata, and analytical samples**
Cohort-wide imaging availability, exact task-specific counts, outcome fields, native TR/acquisition availability and the actual follow-up clinical reference protocols. Main-table sample counts are not the denominator for every task. PCA_protocols lists the imaging dimension budget and selection rule for each analysis.

Numerical results: Dataset S1.

**Table S2. Configurations, checkpoints, inference families, and coverage**
Model input and readout specifications, checkpoint identifiers, correction-family definitions, consistency summaries and analytical coverage. Each result remains associated with its own reference, evaluation protocol and correction family.

Numerical results: Dataset S2.

**Table S3. Atlas- and input-matched FC and FC-Tucker comparisons**
Separate atlas-matched and input-window-matched comparisons. Whole-scan Schaefer100 inputs are not duration-matched to the fMRI models. FC-Tucker is unavailable for Schaefer100 because time series were not supplied. Frozen/adapted scores must be read against their own paired reference.

Numerical results: Dataset S3.

**Table S4. Final-layer readout comparisons**
Current final-normalized-layer alternatives, matched FC/FC-Tucker/input-FC comparisons, dimension-matched identification, cognitive-axis transfer and intervention readouts. Earlier primary-release readouts are labeled earlier; their P values are not interchangeable with the completed final-layer extension. Training-partition readout selection adds 164 paired contrasts and all 540 outer-fold selections in S4_selected_readout_tests and S4_selected_readout_folds. These new analyses retain their own families and no additional absolute-null claims.

Numerical results: Dataset S4.

**Table S5. Frozen prediction and cross-visit identification**
Primary frozen predictions, diagnostic covariate checks and identification. Absolute identity tests retain full readout/projection correction families; their primary-only worksheet is a view, not a new correction. All-cause and AD-etiology dementia are distinct NACC tasks. HRV-ER age category is not age in years. The S5_HRV_age_category worksheet contains the complete eleven-method baseline and follow-up age-category comparison, including the two FC-based reference methods.

Numerical results: Dataset S5.

**Table S6. Fixed-source cognitive transfer and intervention LOSO**
Fixed-source axes are distinct from target-local probes. MEM LOSO uses four intervention cohorts; EF uses five. HRV-ER delayed recall is a one-way external memory endpoint, not a memory-change score or a training cohort in the other MEM LOSO rows.

Numerical results: Dataset S6.

**Table S7. Fusion, covariates, input context, and PCA sensitivity**
Dimension-controlled fusion, covariate/weighting sensitivities, native-80-frame and complete-context analyses, and expanded paired-visit PCA selection. Detailed fold-level PCA selections are supplied in Dataset S14. Sensitivities change one defined factor; they are not cumulatively optimized pipelines. The local HRV-ER delayed-recall fixed5/selected-PC scores and comparisons are provided alongside clearly labeled historical 100-PC results.

Numerical results: Dataset S7.

**Table S8. Paired-visit intervention analyses**
All primary paired-visit cognitive-change and arm-discrimination results, including frozen/post-trained states, baseline/change/combined imaging and clinical increment. No HRV-ER memory-change endpoint exists. Adapted predictions average seeds; comparison counts are not independent experiments.

Numerical results: Dataset S8.

**Table S9. Physiological analyses**

Independent-window nine-configuration physiology, earlier contextual protocols and historical nulls. Usability-filtered exports are historical only. Heart/pulse rate is distinct from variability measures. Absolute physiological tests of the complete covariate-plus-embedding procedure are provided separately in Table S12.

Numerical results: Dataset S9.

**Table S10. Supervised and self-supervised post-training**

Within-source fold-specific encoder evaluation, external cognition, age/sex after SSL, diagnosis, LOSO, identification and clinical increments, plus 162 external and 540 CV checkpoint records. Focused external readout results are complete; the focused adapted-readout identification extension is outside this release. Primary-readout post-training identification is already available.

Numerical results: Dataset S10.

**Table S11. Embedding alignment**

Mean/SD and CORAL alignment for fixed-source cognition, local follow-up/recall and paired-visit tasks. Same-PCA unaligned controls, archived primary controls and Schaefer100 FC are separate contrasts. Exact alignment/standardization invariances retain P=1.

Numerical results: Dataset S11.

**Table S12. Absolute tests and evidence-matrix support**

Absolute tests of primary frozen representations, native-dimensional top-1 identity tests, matched imaging-only follow-up comparisons, numerical reproduction checks and evidence supporting Table S14. Predictive and physiological tests used 1,999 refitted permutations or circular shifts; identity tests used 20,000 mappings. Absolute support, superiority over FC and cross-cohort consistency are separate criteria.

Numerical results: Dataset S12.

**Table S13. Re-ID controls, temporal-order sensitivity, and motion analyses**

Nine-configuration pretrained/random and signal controls, synchronous temporal shuffling, and seven-cohort FD-adjusted Re-ID. S13_FD_ReID_tests contains 3,024 current motion contrasts; S13_nine_config_controls retains 2,448 earlier control rows, including 432 historical pSOPT-only motion contrasts. Only the motion inference is superseded by the expanded correction. S13_temporal_order retains 504 contrasts. The separate scan-average FD experiment supplies 112 fits and 133 paired comparisons. Complete scores, uncertainty, correction families and definitions are supplied externally; no participant-level predictions or large numerical tables are embedded in this document.

Numerical results: Dataset S13.

### Table S14. Evidence for primary frozen representations across tasks

| Configuration | Cross-sectional | | | | | Intervention | | | | | Re-ID |
|---|---|---|---|---|---|---|---|---|---|---|---|
| | Age | Sex | Diagnosis | Memory | EF | Follow-up | | Change | | Arm | |
| | | | | | | Memory | EF | Memory | EF | | |
| BrainLM 13M | ● ○ | ○ ● | ● ● | ● ● | ● ○ | ○ ○ ○ ○ ○ | ○ ○ ○ ○ ○ | ○ ○ ○ ○ – | ○ ○ ○ ○ ○ | ○ ○ ○ ○ ○ | ○ ○<br>○ ● ○ ● ○ |
| BrainLM 111M | ● ○ | ● ● | ● ● | ● ● | ● ○ | ○ ○ ○ ○ ○ | ○ ○ ○ ○ ○ | ○ ○ ○ ○ – | ○ ○ ○ ○ ○ | ○ ○ ○ ○ ○ | ● ○<br>○ ● ● ● ○ |
| BrainLM 650M | ● ○ | ● ● | ● ● | ● ● | ● ○ | ○ ○ ○ ○ ○ | ○ ○ ○ ○ ○ | ○ ○ ○ ○ – | ○ ○ ○ ○ ○ | ○ ○ ○ ○ ○ | ● ●<br>○ ○ ● ○ ○ |
| Brain-JEPA | ○ ○ | ○ ○ | ○ ● | ○ ● | ○ ○ | ○ ○ ○ ○ ○ | ○ ○ ○ ○ ○ | ○ ○ ○ ○ – | ○ ○ ○ ○ ○ | ○ ○ ○ ○ ○ | ○ ○<br>○ ○ ○ ○ ○ |
| Brain-Semantoks | ● ● | ● ● | ● ● | ● ● | ● ○ | ○ ○ ○ ○ ○ | ○ ○ ○ ○ ○ | ○ ○ ○ ○ – | ○ ○ ○ ○ ○ | ○ ○ ○ ○ ○ | ● ●<br>● ○ ● ● ● |
| fMRI-LM VQ-contrastive | ○ ○ | ○ ● | ○ ● | ○ ● | ○ ○ | ○ ○ ○ ○ ○ | ○ ○ ○ ○ ○ | ○ ○ ○ ○ – | ○ ○ ○ ○ ○ | ○ ○ ○ ○ ○ | ○ ●<br>○ ○ ○ ● ● |
| fMRI-LM MAE | ○ ○ | ○ ● | ○ ● | ○ ● | ○ ○ | ○ ○ ○ ○ ○ | ○ ○ ○ ○ ○ | ○ ○ ○ ○ – | ○ ○ ○ ○ ○ | ○ ○ ○ ○ ○ | ○ ●<br>○ ○ ○ ● ● |
| fMRI-LM VQ-domain | ○ ○ | ○ ● | ○ ● | ○ ● | ○ ○ | ○ ○ ○ ○ ○ | ○ ○ ○ ○ ○ | ○ ○ ○ ○ – | ○ ○ ○ ○ ○ | ○ ○ ○ ○ ○ | ○ ●<br>○ ○ ○ ● ○ |
| BrainHarmonix-F | ○ ○ | ○ ● | ● ● | ● ● | ● ○ | ○ ○ ○ ○ ○ | ○ ○ ○ ○ ○ | ○ ○ ○ ○ – | ○ ○ ○ ○ ○ | ○ ○ ○ ○ ○ | ● ○<br>○ ○ ○ ○ ○ |

Order from left to right: cross-sectional, ADNI / NACC; intervention, ACT / CogTE / BREATHE / pSOPT / HRV-ER. Re-ID: ADNI / NACC on the upper line and the five intervention cohorts on the lower line. ●, corrected above-chance performance; ○, no corrected above-chance evidence; –, outcome not available; EF, executive function. Only Brain-Semantoks exceeded its designated FC reference after correction, for follow-up memory in ACT. Full comparisons are in Table S12.

### Table S15. Regional signal variability controls for cognitive and intervention outcomes

Complete SD, primary-readout, embedding-connectivity and FC comparisons across seven cohorts. Results cover cognitive levels, baseline-imaging follow-up prediction, and baseline/change/combined inputs for cognitive-change associations and arm classification, including clinical increments where applicable. All 2,696 paired comparisons and correction families are supplied externally; no large numerical table is embedded here. Re-ID SD comparisons remain in Table S13.

Numerical results: Dataset S16.

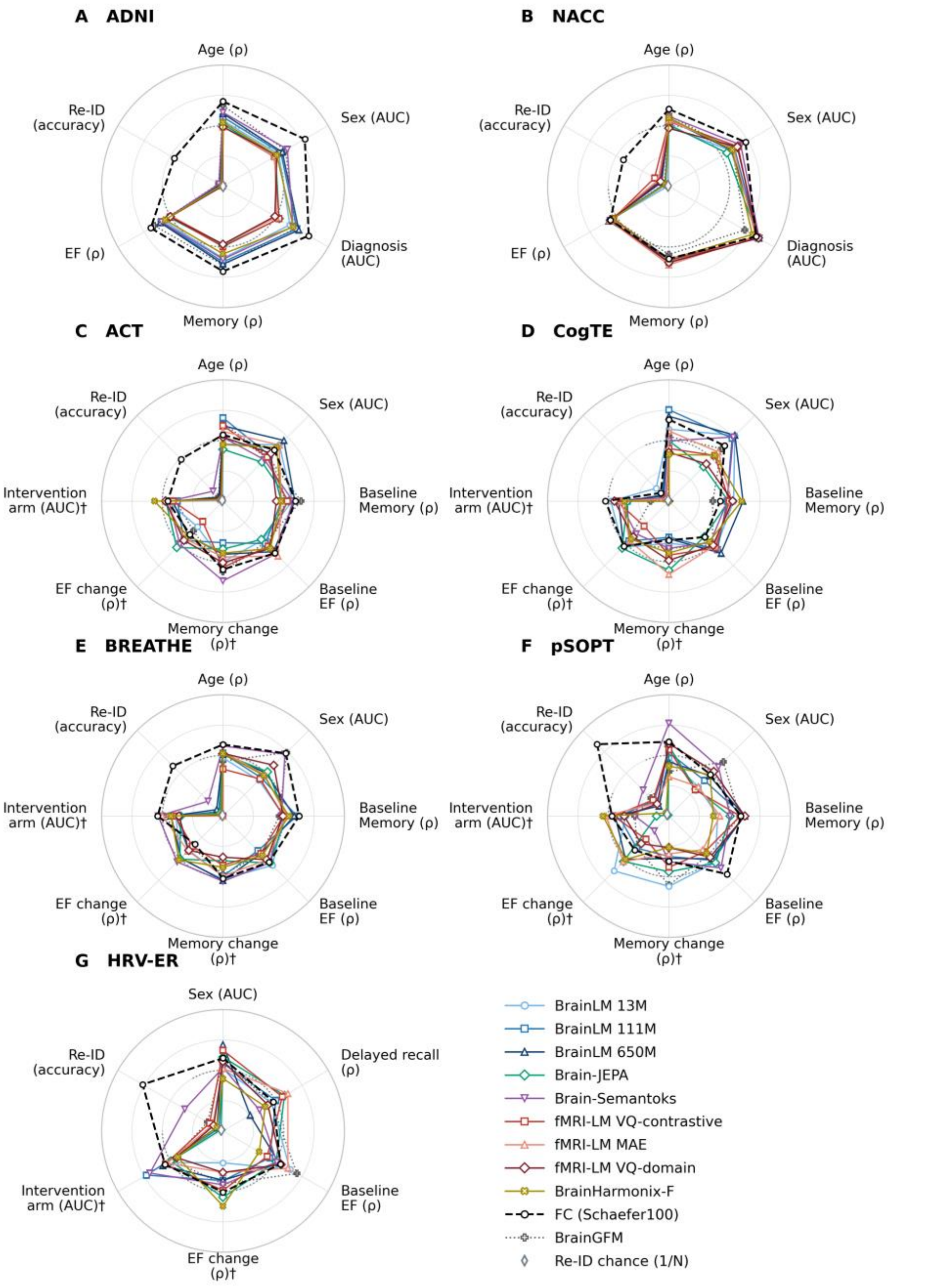


Fig. S1. Predictive performance and cross-visit re-identification across seven aging cohorts. All encoders are frozen and use primary readouts; predictive tasks use imaging only. Cognitive-change and intervention-arm tasks (†) use follow-up-minus-baseline representations. Other predictive tasks use baseline or cross-sectional MRI; HRV-ER delayed recall is a single-assessment outcome predicted from baseline MRI. Re-ID uses paired visits and bidirectional top-1 accuracy. Samples are matched across methods within each task, but can differ between tasks. Radial positions are (ρ + 1)/2 for correlations, AUC for classification, and accuracy on a 0–1 scale for Re-ID. The dotted arc denotes ρ = 0 or AUC = 0.5; the open gray diamond on each Re-ID spoke marks chance (1/N), where N is the number of participants in the gallery. Multiclass AUC is macro-averaged one-versus-rest. Polygon areas are not aggregate performance scores, and the profiles do not indicate statistical significance. Re-ID, cross-visit re-identification; EF, executive function.

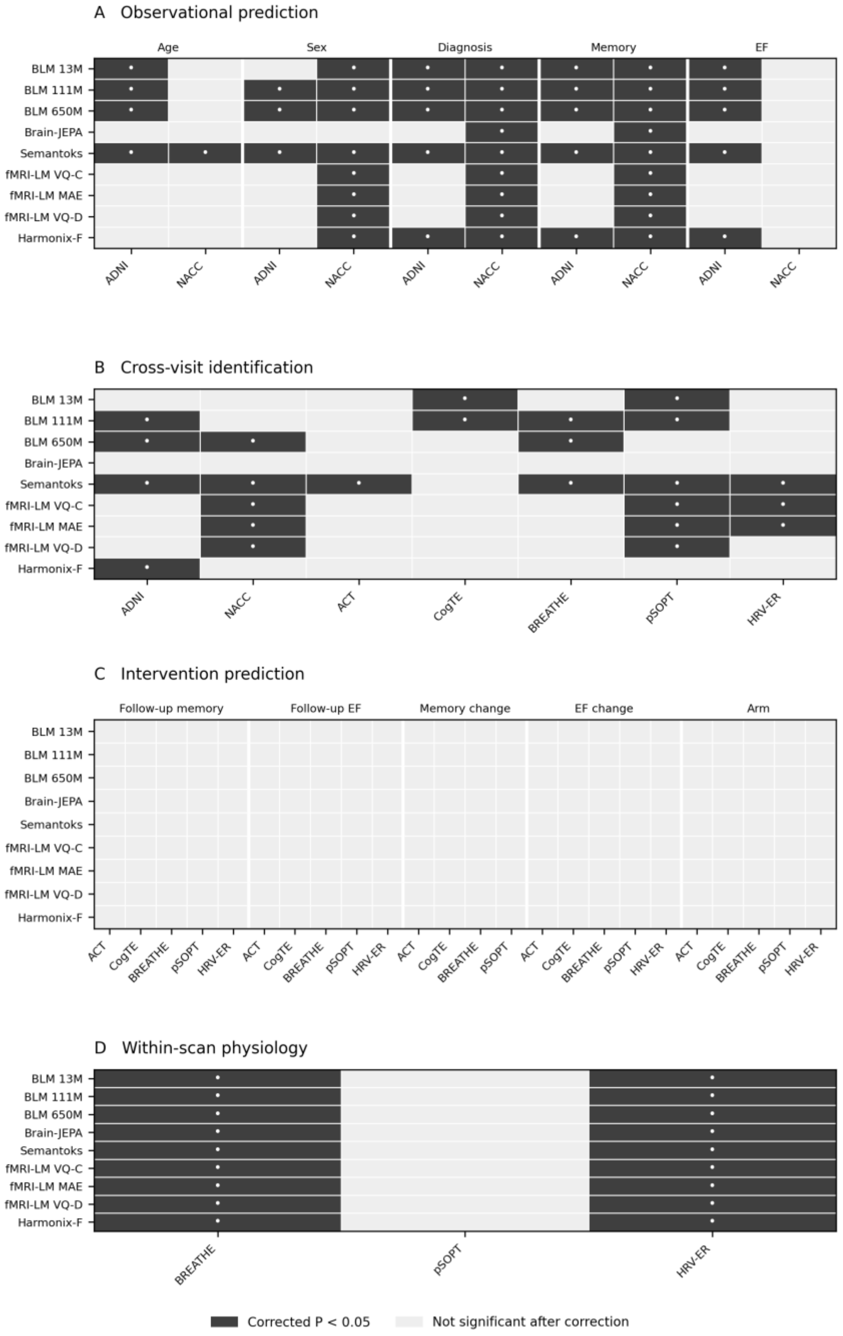


Fig. S2. Corrected absolute-test evidence for primary frozen representations. (A) Within-cohort demographic prediction, diagnostic classification, and concurrent cognitive associations in ADNI and NACC. (B) Primary-readout bidirectional top-1 identification. (C) Imaging-only follow-up cognitive prediction, cognitive-change associations, and intervention-arm classification; the latter two use follow-up-minus-baseline representations. HRV-ER follow-up memory denotes single delayed recall; no memory-change outcome is available. (D) Temporal alignment of the complete nuisance-plus-embedding within-scan physiological prediction procedure. Dark cells denote one-sided Holm-adjusted $P<0.05$; pale cells denote no corrected rejection and do not establish equivalence. Panels A and C used 1,999 refitted label permutations, with a separate family for each task across eligible cohorts and all nine configurations. Panel B used 20,000 identity mappings; its 175-test family additionally retained historical alternative readouts not displayed here. Panel D used 1,999 scan-wise circular shifts and a 27-test family. Physiological significance does not establish incremental embedding value beyond nuisance terms or FC. These are absolute tests, not paired superiority tests or comparisons of effect sizes. All scores, P values, denominators and scope definitions are in Table S12.

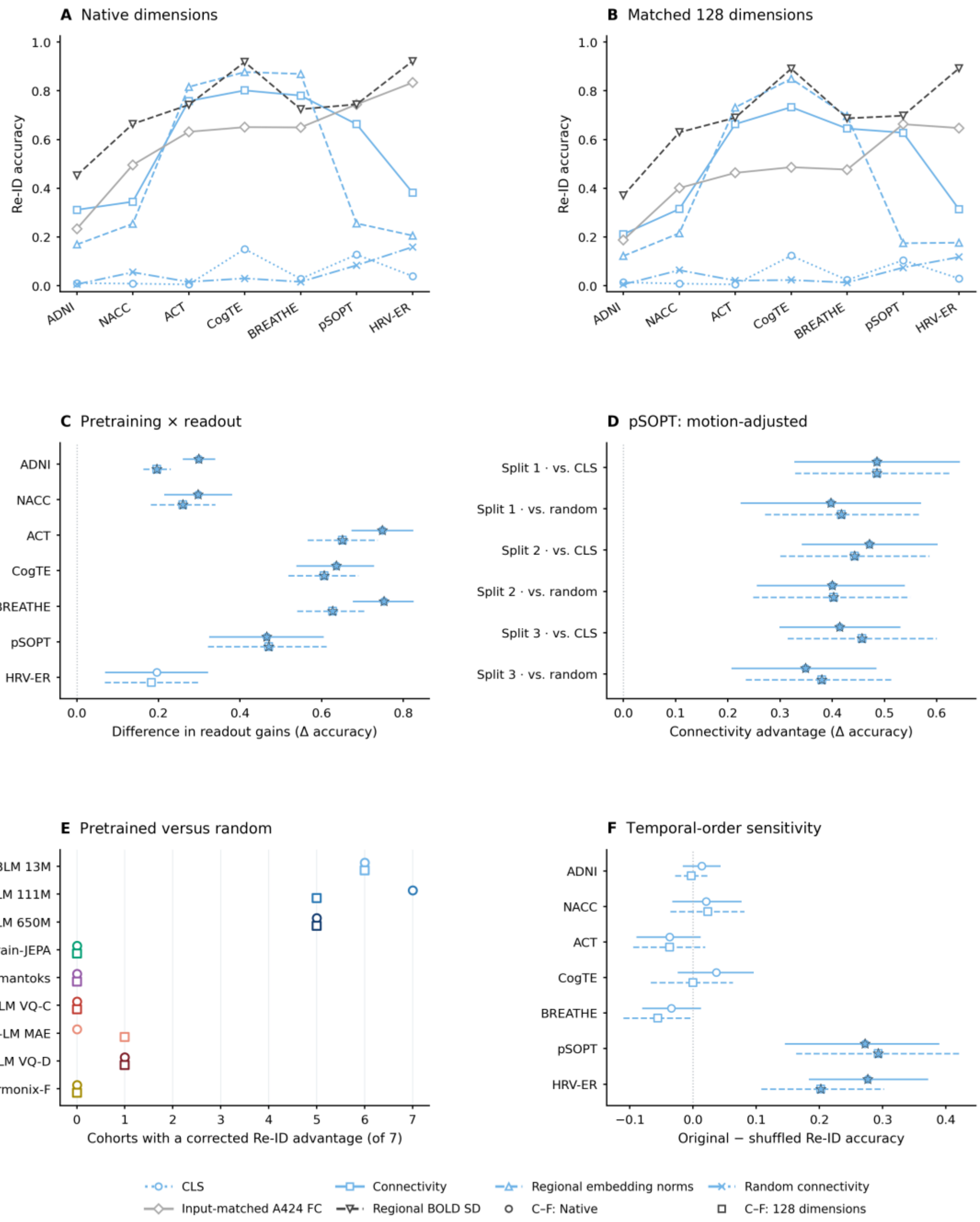


Fig. S3. Readout choice, pretraining, and temporal-order sensitivity in cross-visit Re-ID. (A and B) BrainLM 13M Re-ID accuracy at native dimensionality (A) and after projection to 128 dimensions (B), using final-layer CLS, embedding connectivity, or regional embedding norms, with random-initialization connectivity, input-window-matched A424 FC, and regional BOLD standard deviations as references. Lines connect cohorts for visual comparison. (C) BrainLM 13M pretraining-by-readout interaction: the connectivity-minus-CLS gain with pretrained weights minus the mean gain with random weights. (D) BrainLM 13M connectivity accuracy with original temporal ordering minus the mean accuracy after synchronous time shuffling across regions, which preserved static FC. (E) Number of cohorts in which pretrained connectivity significantly exceeded its random-initialization reference, for each configuration and dimensionality setting. All configurations were evaluated in seven cohorts; these counts are not the benchmark's FC-superiority consistency criterion. (F) BrainLM 13M connectivity advantages over CLS and random connectivity after paired-FD adjustment in pSOPT. Adjustment used separate calibration participants; the three overlapping splits are sensitivity analyses, not independent replications. Random and shuffled reference scores average participant-level scores over five realizations. Circles and squares in C–F distinguish native and 128-dimensional results. Intervals in C, D, and F are pointwise 95% participant-bootstrap intervals conditional on the fixed gallery and realized controls. Panel F uses the seven-cohort FD correction (756 contrasts per metric and dimensionality setting); full FD results across all configurations and cohorts are in Table S13. Stars denote two-sided Holm-adjusted $P < 0.05$ under the applicable complete families; E counts these corrected advantages, while A and B show accuracy without absolute-test annotations. Signal, random, motion, and temporal-order panels retain separate families. Complete results are provided in Tables S4 and S13.

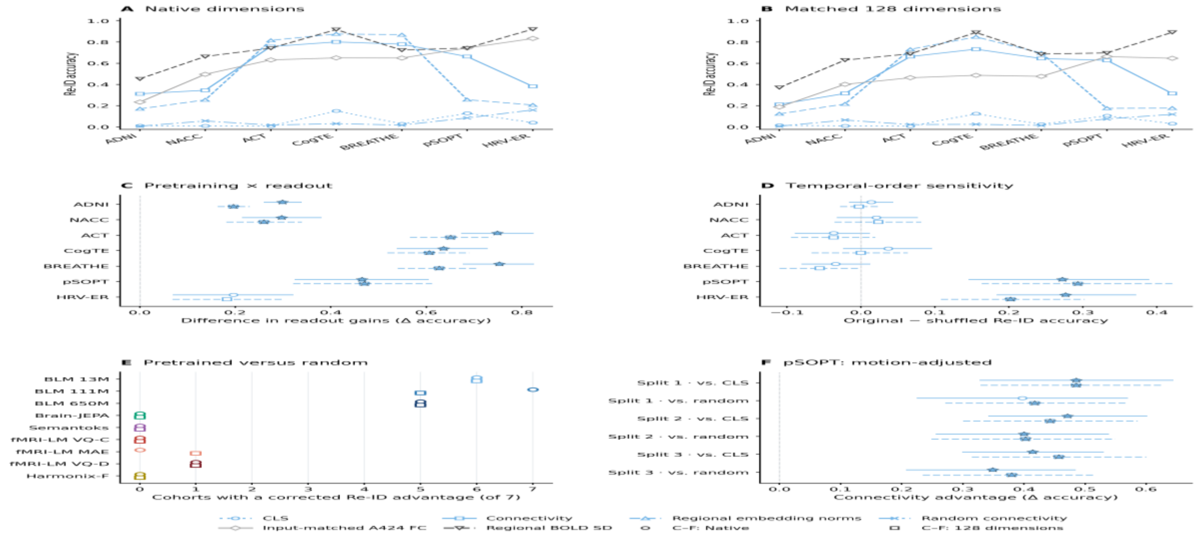


Fig. S4. Individual identity in primary frozen representations. (A) Bidirectional cross-visit identification accuracy for nine time-series FM configurations, Schaefer100 FC and BrainGFM. Gray ticks indicate chance accuracy (1/N) for each cohort-specific gallery. (B) Differential identifiability, defined as mean same-participant minus different-participant similarity. The primary readout is used for each model. Cohorts share participants across the compared methods within each panel; intervention scan pairs span the intervention period. Absolute correspondence-null tests and paired comparisons against FC address different questions and are provided in Table S5. Absolute top-1 coverage includes BrainHarmonix-F under the enlarged 175-test native family; BrainGFM absolute tests and new BrainHarmonix-F differential-identifiability tests are unavailable. Fig. S2 displays corrected primary top-1 support. Points show scores, not independent replications or significance labels.

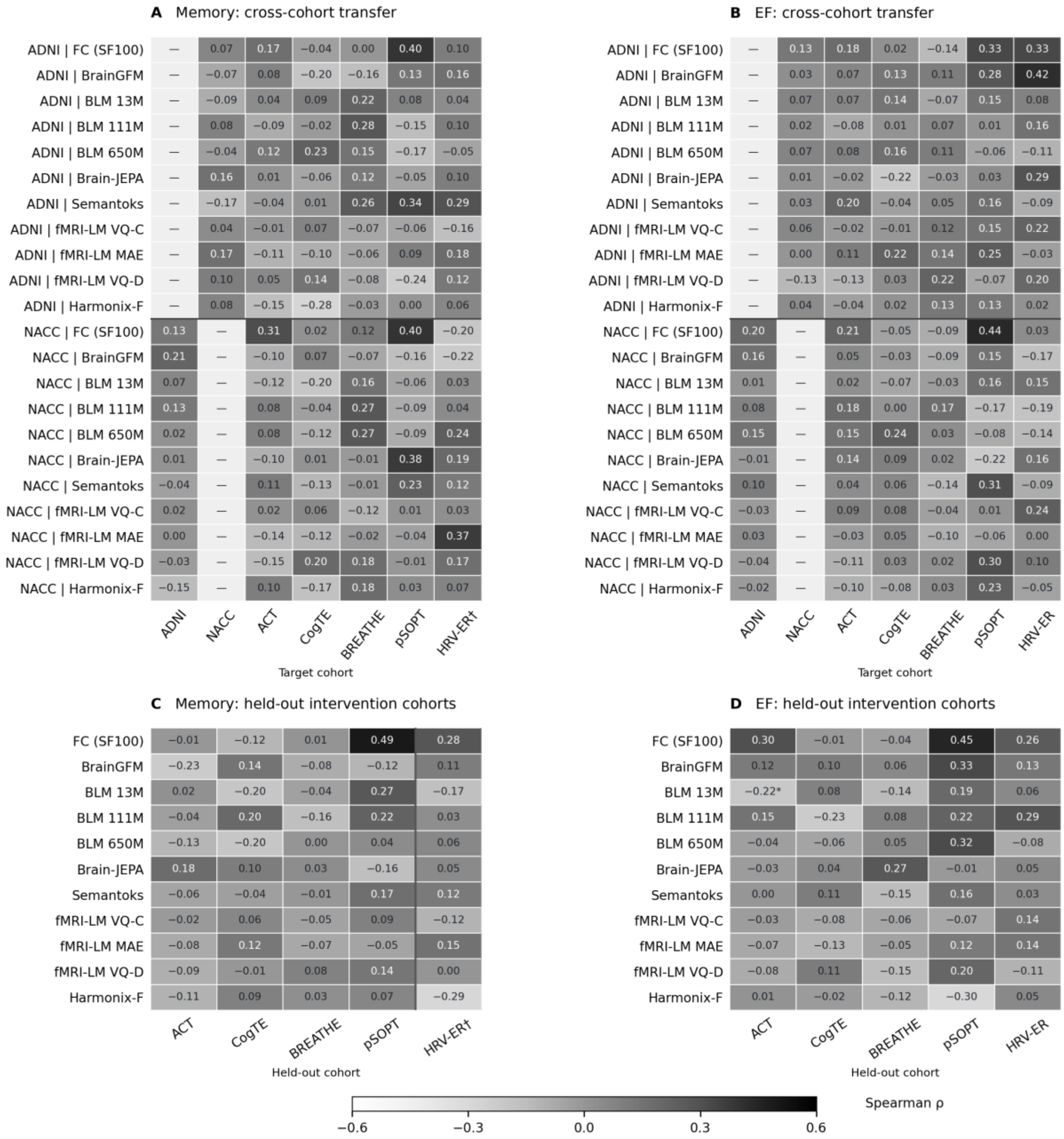


Fig. S5. Cross-cohort transfer of cognitive predictors using frozen representations. (A and B) Memory and EF predictors fitted in ADNI or NACC and applied to target cohorts without refitting. (C and D) Leave-one-study-out prediction across intervention cohorts. HRV-ER memory (†) denotes one-way delayed-recall prediction from the four other intervention cohorts, rather than a fifth memory LOSO fold. Cell values and grayscale intensity indicate Spearman correlation; the continuous color scale does not identify models. A and B are descriptive, without significance inferred from their colors; gray cells denote excluded within-source combinations. Asterisks in C and D indicate Holm-adjusted $P < 0.05$ versus Schaefer100 FC using the complete comparison families; the marked comparison favors FC.

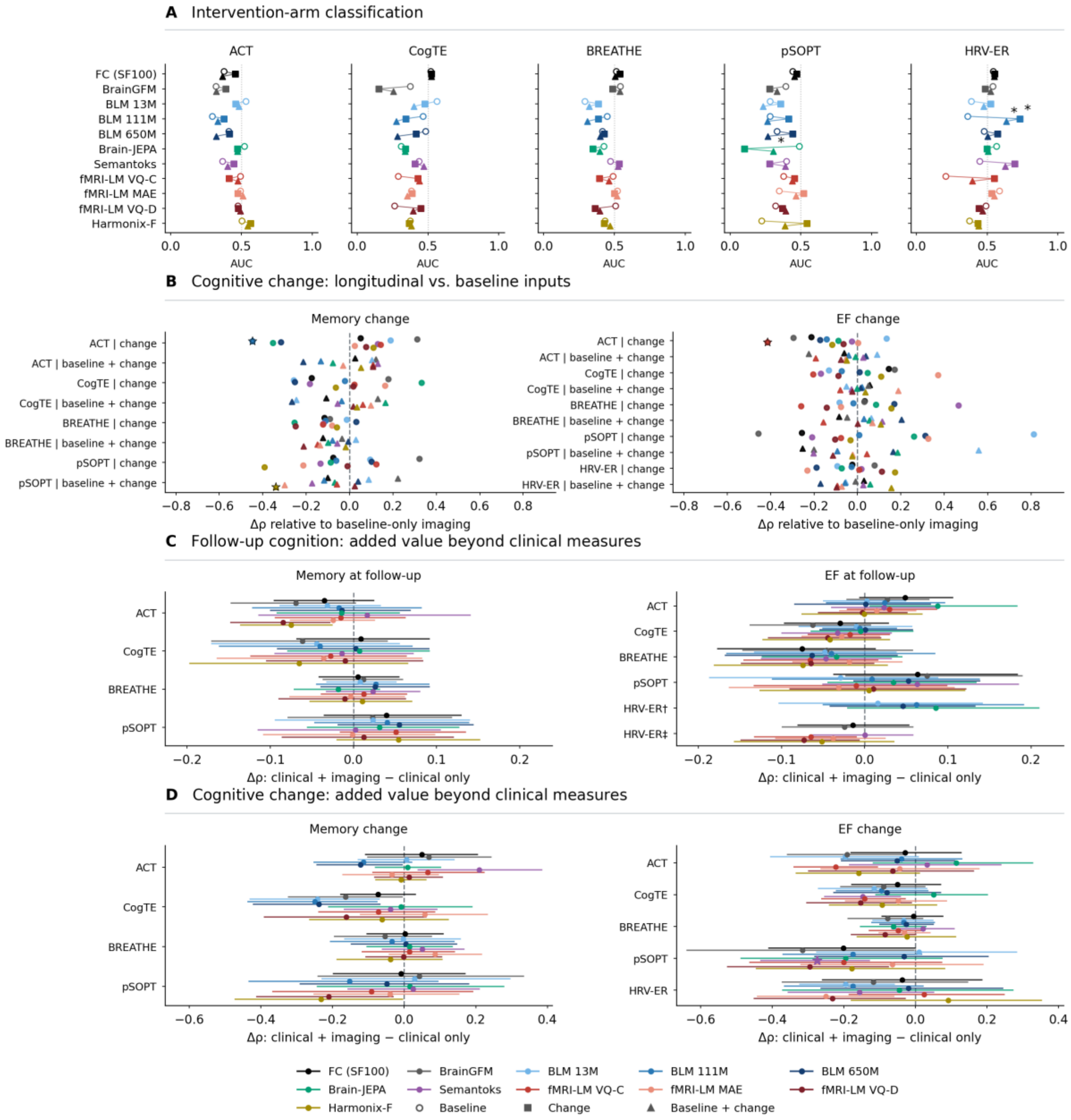


Fig. S6. Longitudinal imaging and incremental predictive value. (A) Intervention-arm classification using baseline representations, change representations (follow-up minus baseline), or their concatenation. Baseline and pre-intervention refer to the same scan. (B) Differences in performance for associations with memory and EF change relative to baseline-only imaging. (C) Incremental value of baseline MRI for predicting follow-up cognitive scores beyond clinical predictors. (D) Incremental value of change representations in analyses of cognitive-change associations beyond clinical predictors. All panels show nine frozen FM configurations, Schaefer100 FC, and BrainGFM. Asterisks indicate Holm-adjusted $P < 0.05$ within the complete corresponding comparison families, including increases and decreases. C and D show pointwise conditional 95% participant-bootstrap confidence intervals. HRV-ER follow-up EF comparisons in C retain two clinical-reference protocols, displayed separately: † the youngest age category and the decrease arm served as reference groups, with indicators for the remaining categories (clinical $\rho = 0.730$); ‡ every age category and both arms had their own indicator, defined within each training partition (clinical $\rho = 0.818$). Both references used baseline EF, age category, sex, and intervention arm in the same 55 participants and cross-validation partitions. Arm-classification sample sizes are ACT 95, CogTE 73, BREATHE 81, pSOPT 43, and HRV-ER 51. B shows point estimates; uncertainty is supplied in the source data. Analyses using follow-up imaging assess longitudinal associations, not pretreatment prediction of response.

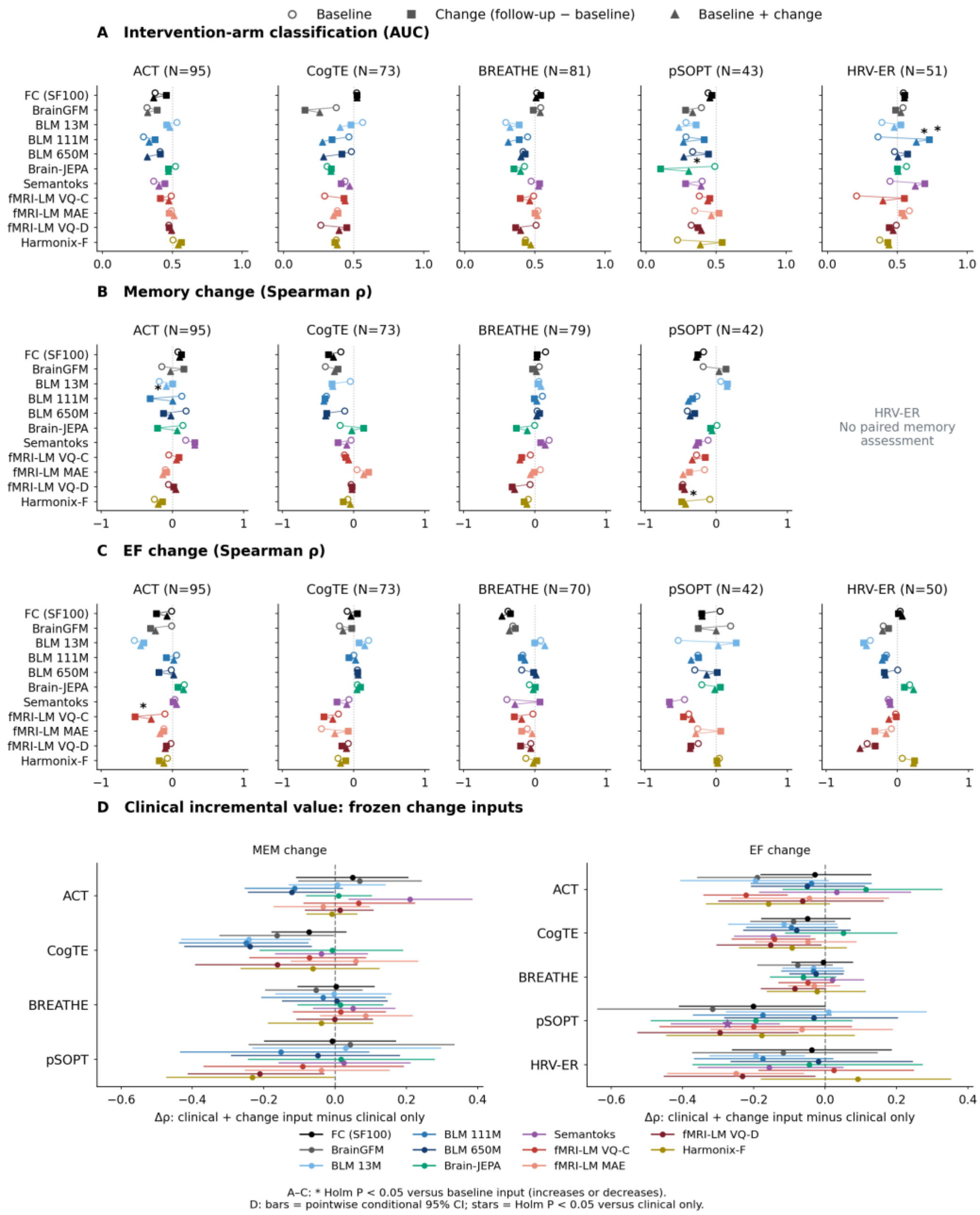


Fig. S7. Complete longitudinal-input comparisons. Intervention-arm classification (A), memory-change associations (B), and EF-change associations (C) use baseline, follow-up-minus-baseline change, or concatenated baseline-plus-change representations from nine frozen FM configurations, Schaefer100 FC, and BrainGFM. Lines connect input choices, not successive visits. Stars indicate Holm-adjusted differences from baseline-only imaging. (D) Incremental value of change representations beyond clinical predictors. Bars indicate pointwise conditional 95% confidence intervals. All significance markers retain the complete comparison families.

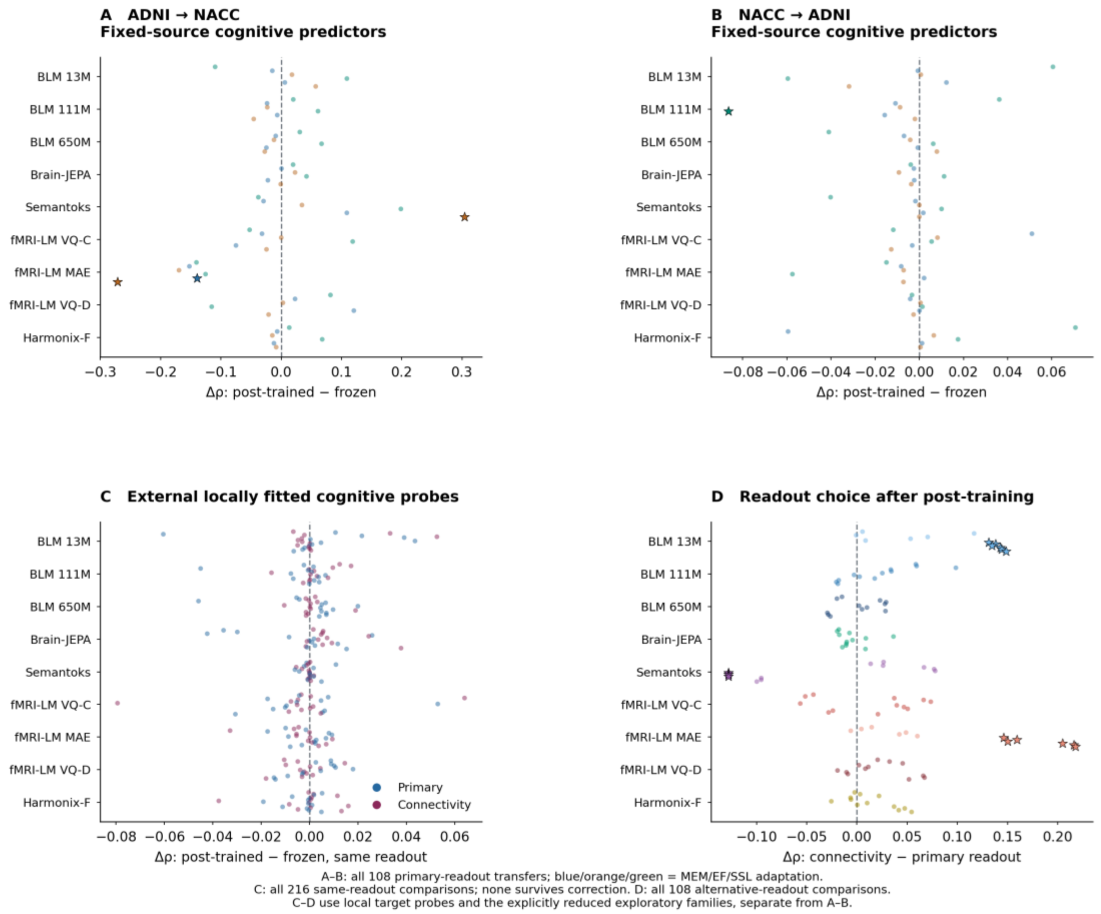


Fig. S8. Complete primary-transfer and focused post-training-by-readout comparisons. Fixed-source cognitive predictors are transferred from ADNI to NACC (A) or NACC to ADNI (B), covering all nine configurations, three adaptation objectives, and both outcomes. (C) All 216 same-readout adapted-versus-frozen comparisons from the focused external concurrent-cognition supplement. (D) All 108 connectivity-versus-primary-readout comparisons after adaptation. C and D use predictors fitted locally in the independent target cohort, unlike A and B. Points represent correlated comparisons; stars indicate Holm-adjusted $P < 0.05$. C and D retain the explicitly reduced exploratory families of the focused supplement. Complete comparisons against Schaefer100 FC and confidence intervals are supplied in Table S10 (Dataset S10).

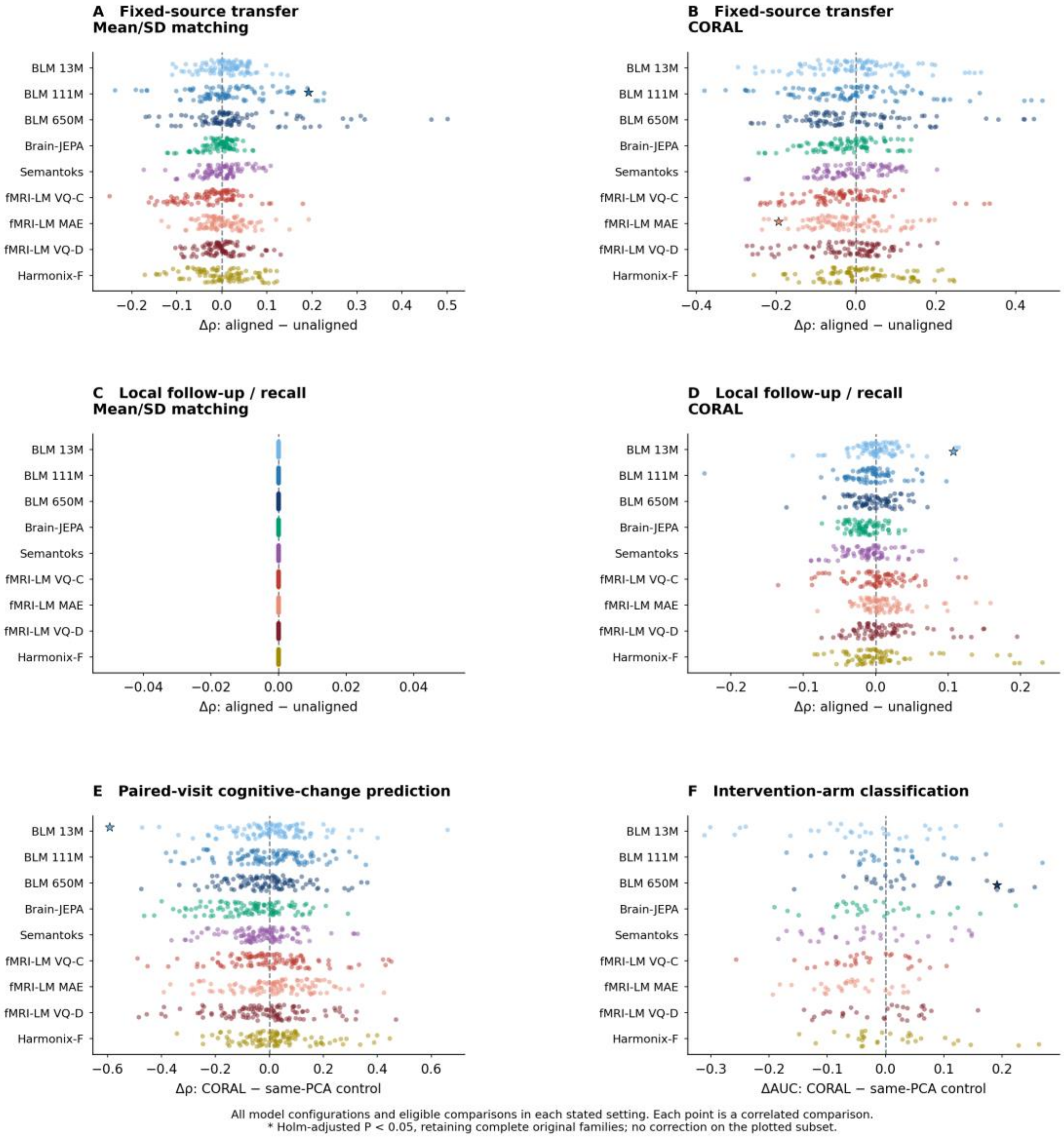


Fig. S9. Embedding alignment across evaluation settings. Mean/standard-deviation matching and CORAL are compared with corresponding unaligned controls for fixed-source cognitive transfer (A and B) and locally fitted follow-up or recall prediction (C and D). Paired-visit CORAL comparisons assess cognitive change (E) and intervention-arm classification (F) against the unaligned control in the same source-fitted PCA coordinates. Every point represents an eligible comparison within the specified setting; points are correlated rather than independent replications. Stars indicate Holm-adjusted $P < 0.05$ within the complete original families. Full comparisons with the primary pipeline and Schaefer100 FC are included in the accompanying source data.

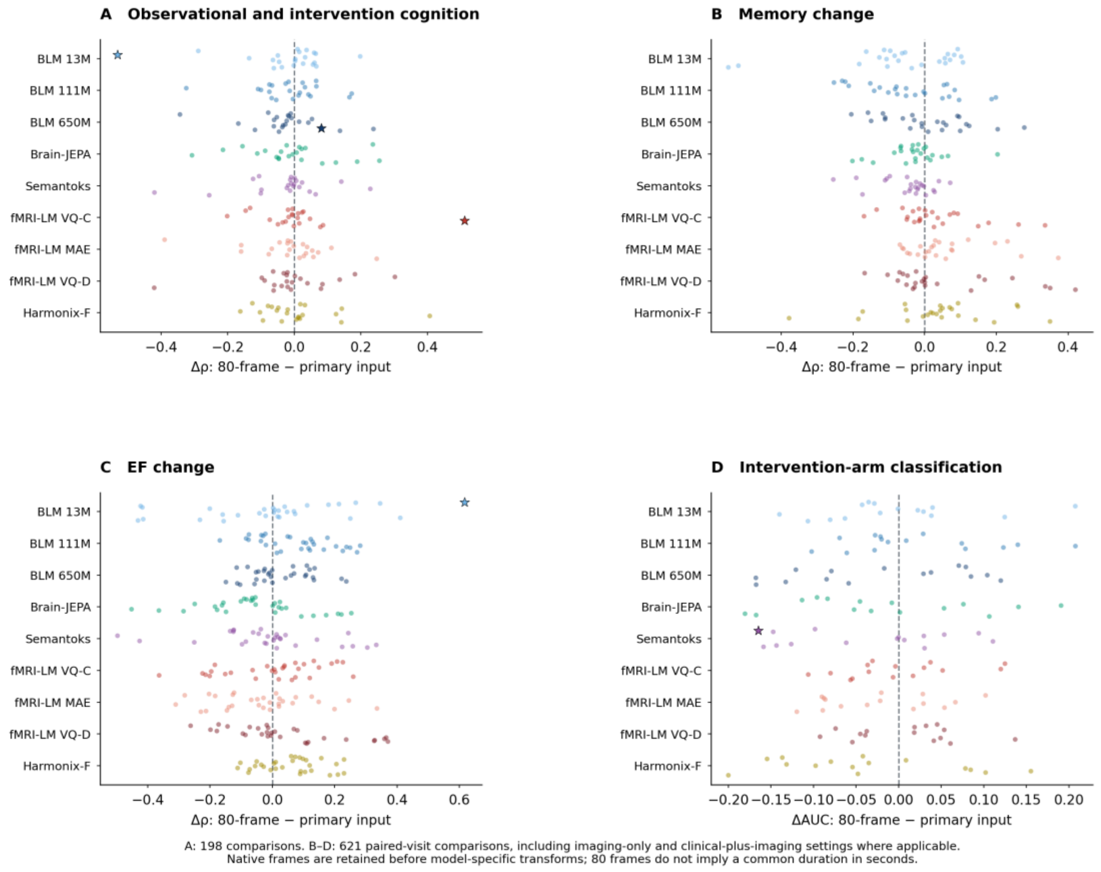


Fig. S10. Sensitivity to input context. Performance using the first 80 native frames is compared with the corresponding primary input for cognitive evaluation in observational and intervention cohorts (A), memory-change associations (B), EF-change associations (C), and intervention-arm classification (D). A includes 198 comparisons; B–D include all 621 short-versus-primary paired-visit comparisons across eligible imaging-only and clinical-plus-imaging settings. Native frames undergo model-specific input construction; equal frame counts do not imply equal durations across cohorts. Stars indicate Holm-adjusted $P < 0.05$ using the complete original families.

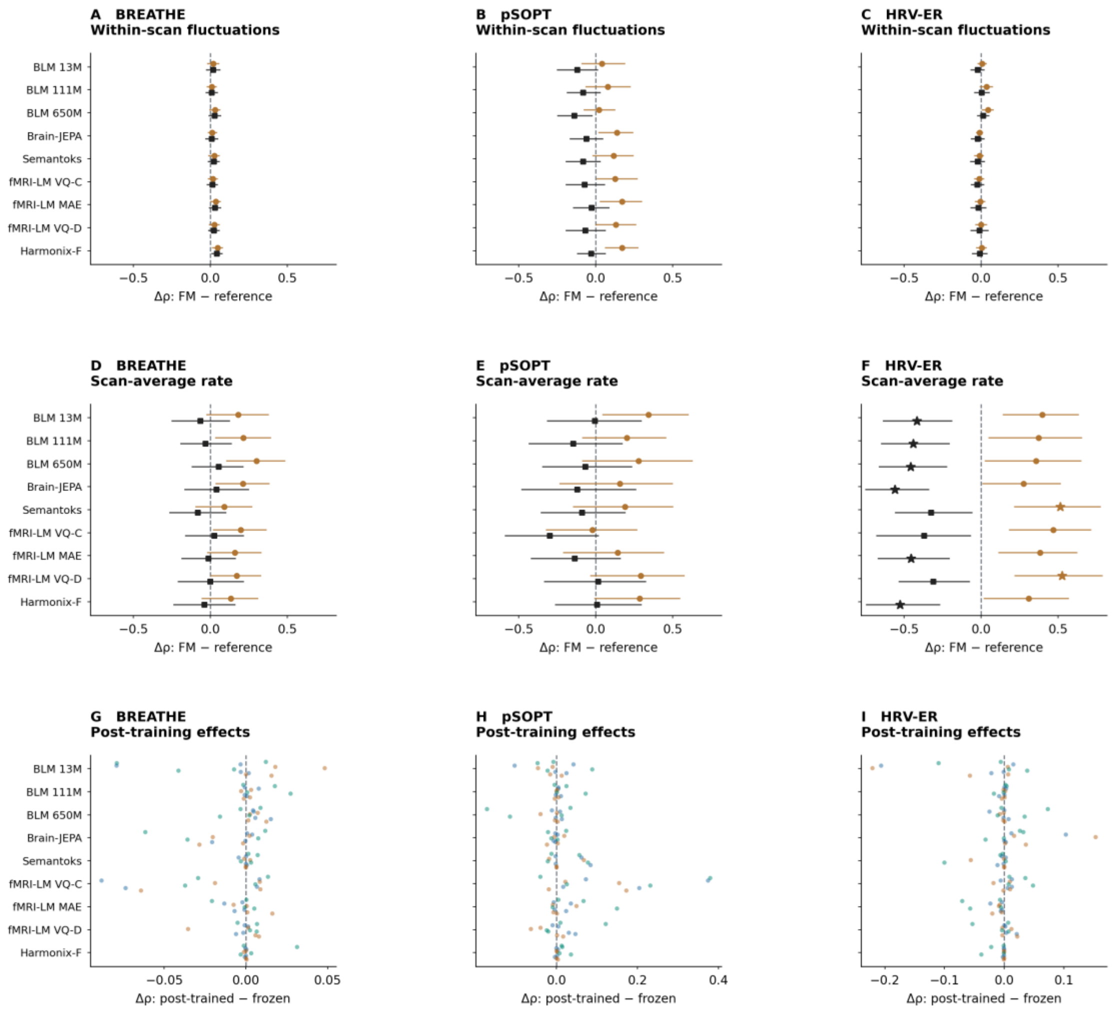


Fig. S11. Physiological prediction using independently encoded windows. Frozen FM representations are compared with covariate-only and atlas-matched FC references for within-scan heart or pulse-rate fluctuations (A–C) and scan-average rate (D–F) in BREATHE, pSOPT, and HRV-ER. Bars denote pointwise conditional 95% confidence intervals. (G–I) All post-trained-versus-frozen comparisons across adaptation sources, objectives, and both analytical scales. Stars indicate Holm-adjusted $P < 0.05$ within the original families. The independently encoded window protocol supports all nine configurations and is distinct from temporal-token pooling with full-scan context. Positive prediction correlations alone are not tests against zero or evidence of superiority to a reference.
